# Shaking hands is a putative terminal selector and controls axon outgrowth of central complex neurons in the insect model *Tribolium*

**DOI:** 10.1101/2020.11.24.393207

**Authors:** Natalia Carolina Garcia-Perez, Gregor Bucher, Marita Buescher

## Abstract

Individual cell types are specified by transcriptional programs which act during development. Gene regulatory mechanisms which specify subtype identity of central complex (CX) neurons are the subject of intense investigation. The CX is a compartment within the brain common to all insect species. The CX functions as a “command center” by initiating motor actions in response to incoming information. The CX is made up of several thousand neurons with more than 60 morphologically distinct identities. Accordingly, transcriptional programs must effect the specification of at least as many neuronal subtypes. Here we demonstrate a role for the transcription factor Shaking hands (Skh) in the specification of embryonic CX neurons in *Tribolium*. The developmental dynamics of *Tc*-*skh* expression are characteristic for terminal selectors of neuronal subtype identity. In the embryonic brain *Tc*-*skh* expression is restricted to a subset of neurons, many of which survive to adulthood and contribute to the mature CX. *Tc*-*skh* expression is maintained throughout the lifetime of the respective CX neurons. *Tc*-*skh* knock-down results in severe axon outgrowth defects thus preventing the formation of an embryonic CX primordium. The as yet unstudied *Drosophila skh* shows a similar embryonic expression pattern suggesting that subtype specification of CX neurons may be conserved.

## Introduction

The insect brain contains a large number of neurons with distinct identities. Cell identity is manifest in specific structural and functional features which together define a neuronal subtype. Subtype identity is determined in the early postmitotic neuron and during development it effects proper axon pathfinding thus facilitating the formation of specific neural connections. Neuronal subtypes express distinct sets of differentiation genes which together bring about all the characteristic features of the cell. Transcription factors that regulate the expression of differentiation genes are the endpoint of hierarchical gene regulatory cascades that act earlier during development. (Hobert, 2008; Hobert, 2011; Allan and Thor, 2015; Hobert and Kratsios, 2019). The early regulatory cascades which govern neuronal subtype specification have been intensively investigated in the insect model *Drosophila melanogaster*, reviewed in (Skeath and Thor, 2003; Lin and Lee, 2012; Crews, 2019). All cells of the *Drosophila* brain derive from embryonically born stem cells, called neuroblasts (NBs). Each NB gives rise to a stereotyped and invariant lineage of neurons and glia. Each NB has a unique identity that is manifested in the expression of a unique combination of transcription factors (Urbach and Technau, 2003). NB identity is determined by overlapping spatial information in the procephalic neuroectoderm. Additional neuronal diversity is generated by a temporal cascade: each NB expresses distinct transcription factors in an invariant temporal series. Temporal factors are inherited by the NB progeny and establish neuronal cell fates characteristic for a given temporal window (Kohwi and Doe, 2013; Lin and Lee, 2012; Rossi et al., 2017; Doe, 2017). The expression of temporal transcription factors can be transient, thus making them unlikely regulators of differentiation genes which need to be expressed throughout the life of a neuron (Sullivan et al., 2019). In addition, Notch-signaling generates subtype diversity: sibling neurons take on different cell fates and form hemi-lineages in a Notch-ON/Notch-OFF dependent manner (Buescher et al., 1998; Truman et al., 2010). In the *Drosophila* ventral nerve cord (VNC), spatial and temporal factors converge to activate the expression of transcription factors that function as terminal selectors of neuronal subtype identity: these factors regulate the lifelong expression of effector genes that together bring about the structural and molecular features of the differentiated cell type (Allan and Thor, 2015; Hobert and Kratsios, 2019). A role for terminal selectors in the *Drosophila* brain has not been demonstrated as yet.

Current interest in the specification of subtype identity is focused on neurons whose trajectories build up the central complex (CX) (Boyan and Reichert, 2011; Sullivan et al., 2019; Hartenstein et al., 2020). The CX is a prominent compartment in the center of the brain that is common to all insect species. It functions as a multi-modal information processing center which commands locomotor behaviors in response to visual stimuli (Strauss and Heisenberg, 1993; Pfeiffer and Homberg, 2014; Heinze, 2017; Franconville et al., 2018). Anatomically, the adult CX is an ensemble of interconnected paired and unpaired neuropils (Hanesch et al., 1989; Strausfeld, 1999). CX anatomy is particularly well described in *Drosophila*. The core components of the CX are the protocerebral bridge (PB), the fan-shaped body (FB), the ellipsoid body (EB) and the noduli (NO) (for a schematic depiction of the CX, refer to Figure1E). The PB is located at the dorsoposterior cell body-neuropil interface wedged between the two calyces of the mushroom bodies (MB). The PB consists of 16 glomeruli arranged in the shape of a handlebar with its tips bent to a posterior-ventral position. The FB is located anterior-ventrally and forms the largest neuropil of the CX. Within the FB, neuronal trajectories are organized to form an intricate substructure of horizontal strata and vertical slices. Just anteriorly to the FB lies the EB, a ring-shaped neuropil that is structured into radial sectors and concentric zones. While the PB, the FB and the EB are midline spanning neuropils, the ventral most module of the CX, the NO, are paired. Two further, paired, modules are closely associated with the CX: the bulbs (BU) and the lateral accessory lobes (LAL).

**Figure 1.**
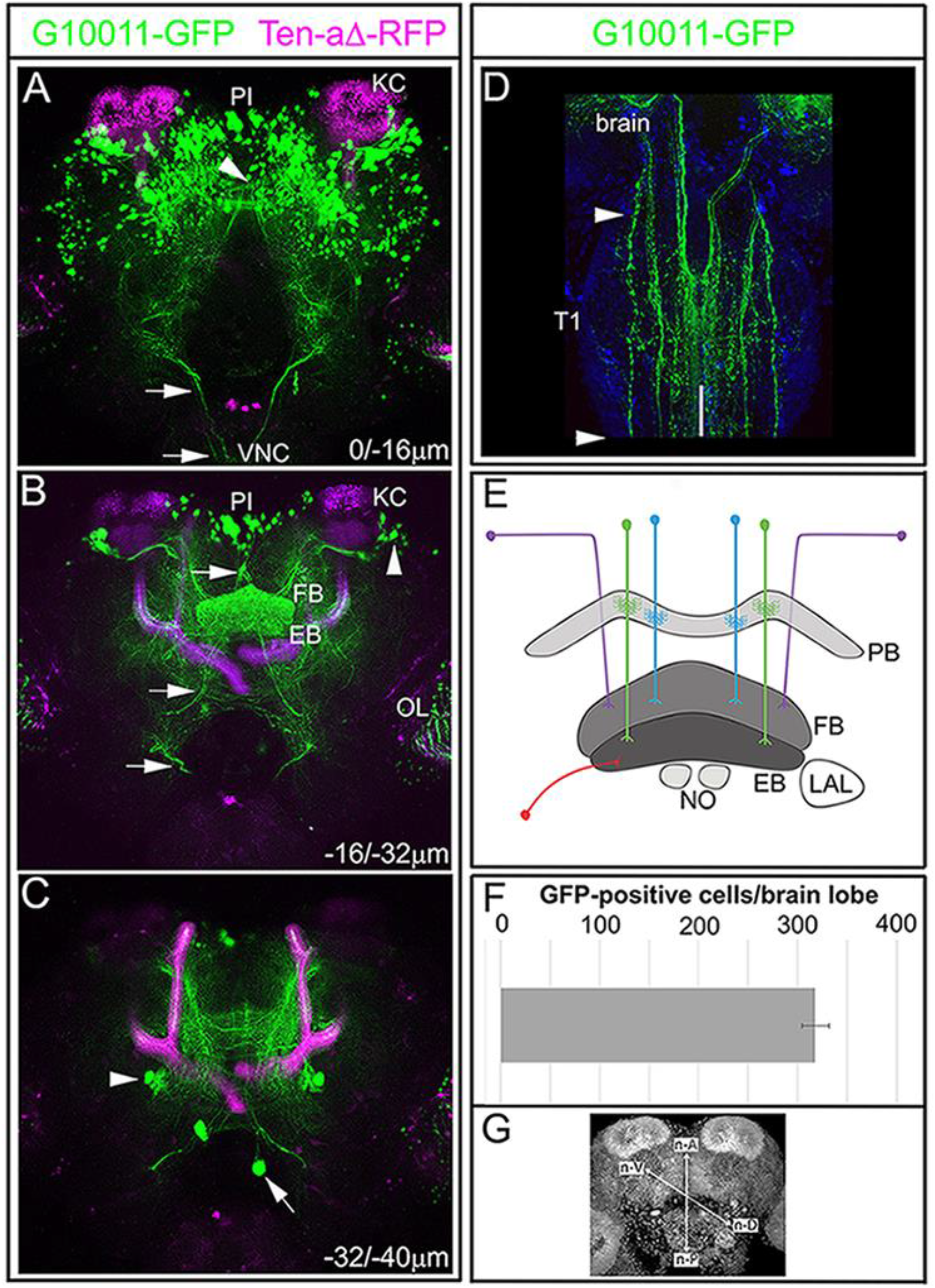
The enhancer trap line G10011-GFP labels a subset of CX neuropils in the adult *Tribolium* brain. Brain of an animal with the genotype G10011-GFP;Ten-aΔ-RFP (GFP auto-fluorescence green, and RFP auto-fluorescence magenta). Note that Ten-aΔ-RFP expression is restricted to the mushroom bodies (MB; magenta). Serial confocal sections were combined and visualized as maximum intensity projections to display distinct anatomical features. Scan direction is from the n-dorsal (n-D) to the n-ventral (n-V) surface of the brain (coordinates of the neuraxes are shown in (**G**). Depth along the Z-axis is given in μm. (**A**) GFP-positive cell bodies in the posterior brain. GFP-expression is absent from the Kenyon cells (KC) of the MB. Arrowhead indicates the protocerebral bridge (PB) which is only partially visible (for a clearer view of the PB, refer to Figure 2 A). Arrows indicate descending axon projections which extend longitudinal connectives into the ventral nerve cord (VNC). (**B**) The fan-shaped body (FB) and the ellipsoid body (EB) are heavily labelled by GFP. Clusters of laterally located cells send their axon trajectories toward the upper layer of the FB body (arrowheads; compare with Figure 2 L). Small clusters of large cells are located in the pars intercerebralis (PI). Based on location and axon projections, they are likely to be neurosecretory cells. GFP-fluorescence is also seen in the optic lobes (OL). (**C**) The arrowhead marks a single cell cluster of 4-6 GFP-positive cells in the anterior brain (arrowhead). Individual large cells within the Tritocerebrum project anteriorally towards the PI and are likely to be homologous to the “hugin expressing” cells which have been identified in several other insect species (arrow). (**D**) First thoracic segment (T1) of the VNC: multiple axon projections which originate in the brain form longitudinal connectives in the VNC (arrowheads mark the limits of the first thoracic segment T1). Note the absence of GFP-positive somata in the VNC. White line marks the ventral midline. (**E**) Schematic illustration of CX small-field and large-field neurons. Two types of small-field neurons, pb-fb-eb (green) and pb-fb (blue) are shown. Two types of large-field neurons, a ring neuron (red) and an AOTU neuron (purple) are shown. (**F**) The average number of G10011-GFP-positive cells in 2-3 day old animals is 320 per brain lobe (n=4). **(G)** Coordinates according to the neuraxes.

Neurons whose projections make up the neuropils of the CX are classified as either small-field or large-field neurons (Young and Armstrong, 2010; Yang et al., 2013; Wolff et al., 2015). The best studied group of small-field neurons are the columnar neurons. They form eight sets of isomorphic cells within each brain hemisphere whose somata reside within the pars intercerebralis (PI). The projections of a subgroup of columnar neurons form dendritic tufts giving rise to the 16 glomeruli of the PB. Further anterior-ventrally, columnar neurons project four prominent bilateral pairs of fiber bundles (w, x, y and z tracts). These tracts connect the PB to the FB by an intricate pattern of inter-hemispheric crossings before they extend further anterior-ventrally to establish the columnar structure of the FB. Large-field neurons provide input from other brain areas into the core of the CX. Some large-field neurons project perpendicular to the columnar neuron tracts and effect the horizontal stratification of the FB. For example, such a projection pattern is characteristic for a subset of neurons of the anterior optic tubercle (AOTU): they project first medially and then ventrally to innervate the uppermost stratum of the FB. Another well-studied group of large-field neurons are the ring neurons whose somata reside ventrolaterally to the CX and whose trajectories innervate the EB.

The architecture of the adult CX and its internal connectivity are well documented in many insect species (Loesel et al., 2002). By contrast, little is known about the regulatory mechanisms which specify subtypes of CX neurons. A few studies have addressed the roles of spatial and temporal factors in the specification of *Drosophila* CX neurons. For example, ring neurons arise from within a spot of *engrailed* expressing procephalic neuroectoderm and loss of Engrailed results in the loss of embryonic ring neurons (Bridi et al., 2019). Recently, the temporal transcription factor Eyeless and its target Twin of Eyeless were shown to specify features of a subset of columnar CX neurons (Sullivan et al., 2019).

While the overall architecture of the CX is well conserved across different insect species, the size and shape of its neuropils vary greatly reflecting an evolutionary adaptation to different habitats (Loesel et al., 2002; Strausfeld, 1999; Koniszewski et al., 2016). Moreover, the assembly of individual CX neuropils occurs at different stages of development, a phenomenon referred to as heterochrony (Panov, 1959). We study the regulatory mechanisms that underlie CX development in the red flour beetle *Tribolium castaneum* (He et al., 2019; Farnworth et al., 2020). *Tribolium* is an insect model well suited to the study of gene regulatory pathways: its genome is fully sequenced (Herndon et al., 2020) and *Tribolium* is amenable to genetic manipulation, including enhancer trapping (Trauner et al., 2009). Additionally, “parental RNA interference” (RNAi) is well established as a means to study gene function (Bucher et al., 2002; Schmitt-Engel et al., 2015). General features of embryonic neurogenesis are remarkably well conserved between *Tribolium* and *Drosophil*a (Wheeler, 2003; Biffar and Stollewerk, 2014).

Here we report a role for the transcription factor Tc-Skh, the *Tribolium* ortholog of *C. elegans* Unc-42, in the specification of a subset of CX neurons. The developmental dynamics of *Tc*-*skh* expression are characteristic for terminal selectors of neuronal subtype identity. *Tc*-*skh* is not expressed in neural progenitors or glia but is expressed in neurons of 14 to 16 embryonic lineages. Many of these neurons survive to adulthood and a subset contributes to the adult CX. Expression of *Tc*-*skh* in CX neurons is maintained into adulthood. Notably, *Tc-skh* is absent from neurons which make up other major compartments of the brain, e.g. the mushroom bodies and the antennal lobes. *Tc*-*skh* RNAi results in severe axon outgrowth defects thus preventing the formation of an embryonic CX primordium. In addition, we observe a moderate reduction of *Tc*-*skh* expression. The *Drosophila* ortholog *Dm*-*skh* shows a highly similar expression pattern in the embryo suggesting a conserved role in the specification of CX neurons.

## Results

### The enhancer trap line G10011 labels several neuropils of the *Tribolium* adult CX

To identify genes which play a role in CX development, we screened a collection of enhancer trap lines that express an untagged version of eGFP (Trauner et al., 2009). Analysis of GFP fluorescence in *Tribolium* adult brains led to the identification of the line G10011 in which the CX is heavily labeled (for a 3-D overview see Figure **S1**). G10011 beetles are homozygous viable, fertile and their lifespan is comparable to that of the *Tribolium* wild type strain *SB*. We did not detect obvious differences in the fluorescence patterns of male and female and young and old adult brains. To gain an overview of G10011-GFP expression, we crossed G10011 to Ten-a-Δ-RFP expressing beetles and examined the adult brains of the resulting progeny. Ten-a-Δ-RFP is a derivative of the enhancer trap line Tenascin-a (also called Teneurin-a)-GFP (He et al., 2019).

In Ten-a-Δ adult brains, RFP expression is restricted to the MBs which provide an internal landmark (Figure 1A-C). Double-labeled brains show GFP fluorescence in the PB (arrowhead in Figure 1A), the FB and the EB (Figure 1B; for a schematic depiction of CX neuropils and coordinates of the axes refer to panels E and G, respectively). G10011-GFP positive somata reside nearly exclusively in the n-dorsal brain. The majority of GFP-positive cell bodies are located in the n-antero-medial region where they form several large clusters within the PI and also more n-posterior areas. Small clusters of large cells are located in the anterior most region of the PI (Figure 1A,B). These cells project descending axons which form a chiasma and then extend further posterior to enter the VNC. Based on cell body location and axonal projections, they are likely to be neurosecretory cells. In the dorsolateral brain large clusters of cells reside lateral to the Kenyon cells (KC) of the MBs (Figure 1B, arrowhead). In addition, a small number of GFP-positive cells is scattered throughout the lateral regions of the n-dorsal brain. The ventral cortex of the brain contains only a single GFP-positive cluster comprising 6-8 cells which is located ventrolaterally to the EB (arrowhead in Figure 1C). Finally, a few large cells within the tritocerebrum project towards the PI and are likely to be “hugin expressing” cells, a type of neurosecretory cells identified in several insect species (arrow in Figure 1C) (Melcher and Pankratz, 2005). We determined an average number of 320 GFP-positive cells per brain lobe (n=4; Figure 1F). Notably, G10011-GFP is not expressed in the Kenyon cells of the MBs and the antennal lobes. We do not know whether GFP expression in the optic lobes is attributable to the G10011 insertion since the transformation marker 3xP3 itself directs GFP expression in the optic lobes (Berghammer et al., 1999). The VNC shows no GFP-positive somata but contains multiple GFP-positive longitudinal connectives which originate in the brain (Figure 1D).

To examine GFP-fluorescence in CX neuropils in more detail, G10011 adult brains were stained with α-Synapsin which facilitates the visualization of individual brain neuropils (Figure 2A-D and E-H’).

**Figure 2.**
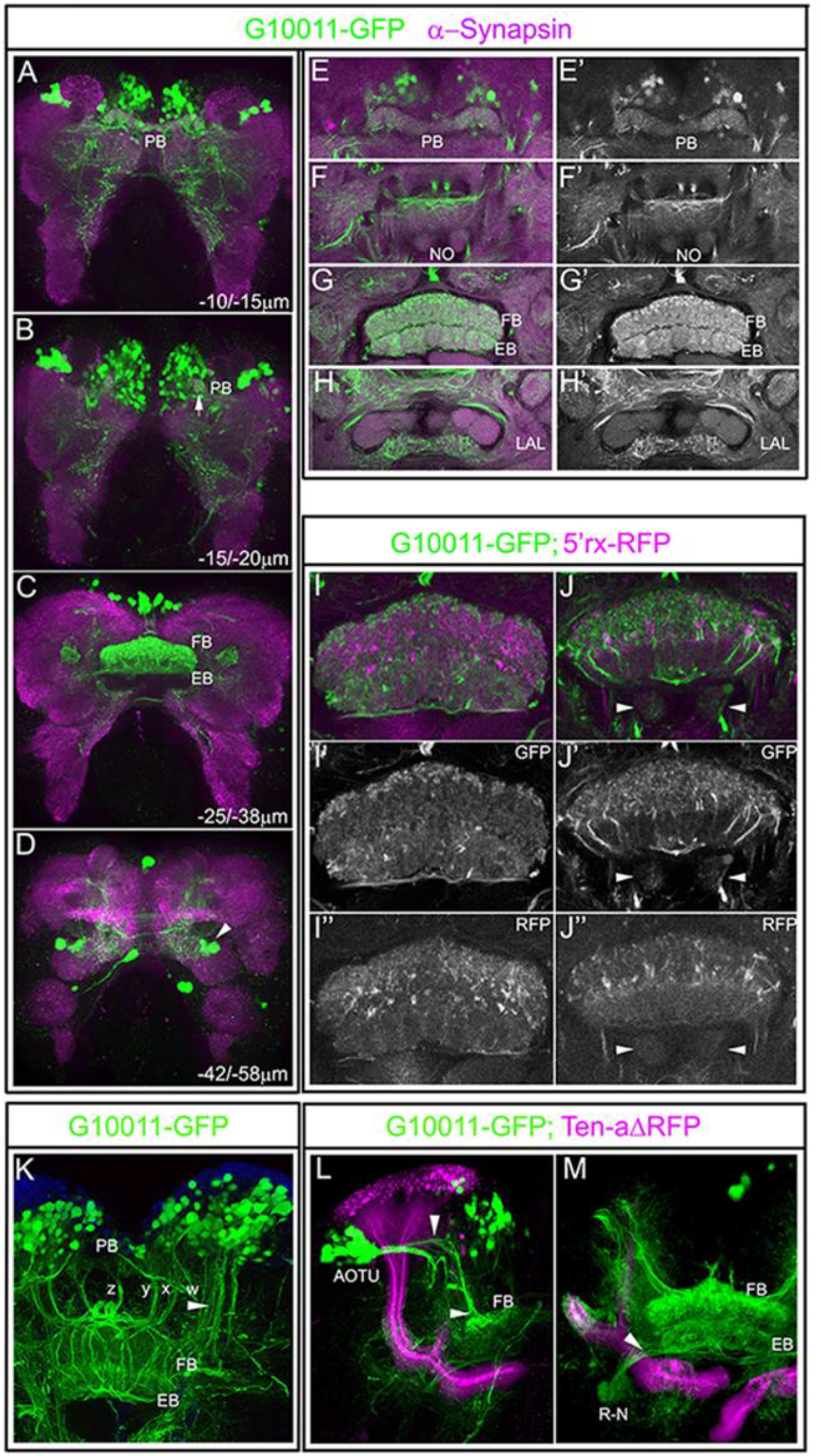
GFP-labeled CX neuropils in the adult G10011 brain. Serial confocal sections were combined and visualized as maximum intensity projections to display individual anatomical features. (**A-H’**) Adult G10011 brains (green, GFP auto-fluorescence) stained with α-Synapsin antibody (magenta). (**A-D**) Whole brain imaged at low magnification. Scan direction is from the n-dorsal to the n-ventral surface of the brain. Depth along the Z-axis is given in μm. (**A-B**) GFP-label within the PB. Arrow in (**B**) indicates the most lateral glomerulus of the PB. (**C**) GFP-label within the FB and the EB. (**D**) GFP-label of a subset of ring neurons (arrowhead). (**E-H’**) Close-ups of CX neuropils. (**E**) PB, (**F**) NO, (**G**) FB and EB, (**H**) We interpret triangular compartments which are located postero-lateral to the EB as the lateral-acessory-lobes (LALs). (**E’-H’**) GFP auto-fluorescence only. Note that the NO and the putative LALs are only weakly labelled by GFP. (**I-J’’**) CX of an animal with the genotype G10011-GFP;5’rx-RFP. (**I-I’’** and **J-J’’**) depict two different planes of the CX along the D/V axis. (**I’, J’**) GFP auto-fluorescence only. (**I’’, J’’**) RFP only. Note that there is little overlap of GFP- and RFP-fluorescence. Arrowheads indicate the NO. (**K-M**) identified sets of neurons with projections into the FB and/or the EB. (**K**) small-field, columnar neurons arborize within the PB, form the characteristic z, y, x and w fascicles, decussate in the upper part of the FB and establish the columnar organization of the FB. Blue stain is DAPI. (**L-M)** Adult brain of an animal with the genotype G10011-GFP;Ten-aΔ-RFP. (**L**) AOTU large-field neurons and their projections into the FB (arrowheads). (**M**) Large-field ring neurons (R-N) projection into the EB body (arrowhead).

Image analysis at both low (Figure 2C) and high magnification (Figure 2G,G’) confirmed that the FB and the EB are strongly labelled by GFP. Within the FB, GFP-fluorescence is observed in all columns and strata with a particularly heavy label of the uppermost stratum. Within the EB, GFP labels all radial segments (Figure 2G,G’). In addition, all 16 glomeruli of the PB are labelled by GFP (Figure 2A,B and E,E’). In contrast to the strong GFP-label within the midline-spanning neuropils, GFP-fluorescence within the paired NO is very weak (Figure 2F,F’). The CX modules are associated with additional neuropils such as the BUs and the LALs. In *Tribolium* both of these compartments have not been described as yet. We observed a bilaterally symmetric brain area located posterior-ventrally to the FB/EB which we interpret as the LALs. G10011-GFP fluorescence within the putative LALs is weak (Figure 2H,H’). We were not able to identify a structure which may represent the BU.

Strong GFP-fluorescence in the FB and the EB raises the question as to whether G10011-GFP labels all neuronal projections that make up these neuropils. To address this question, we crossed G10011 beetles to the imaging line 5’*rx* (*retinal homeobox gene*) in which the FB and the EB are intensely labelled by RFP (He et al., 2019). Image analysis of the respective progeny revealed that G10011-GFP and 5’*rx*-RFP fluorescence are largely non-overlapping, indicating G10011-GFP labels only a subset of structures within the FB and the EB (Figure 2I-J’’).

Anatomical studies in a variety of insects have led to the characterization of small- and large-field neurons whose projections make up the neuropils of the CX (Hanesch et al., 1989; Young and Armstrong, 2010; Yang et al., 2013). Cell body location, morphology and projections of many CX neurons are conserved among different insect species (Pfeiffer and Homberg, 2014). Based on these criteria, we were able to identify one type of small- and two types of large-field neurons that express G10011-GFP. Firstly, sets of neurons whose cell bodies reside within the PI show all properties indicative of columnar neurons (Figure 2K): their trajectories contribute to the dendritic tufts within the glomeruli of the PB and then extend more posterior to form four characteristic fiber tracts commonly named z, y, x and w tracts. These tracts connect the PB to the FB by interhemispheric chiasmata before they extend further posterior to establish the columnar structure of the FB (Figure 1K). Secondly, in the dorsolateral brain two large clusters of neurons adjacent to the MB calyces project two major fiber bundles, one of which extends first medially and then posterior before it enters the uppermost stratum of the FB (Figure 2L). We interpret these CX neurons as a subset of the AOTU neurons. Thirdly, we identified the ring neurons (R-N) whose somata reside posterior-laterally to the EB and whose projections innervate the EB (Figure 2M). There are likely to be more G10011-GFP positive neurons contributing to the neuropils of the CX. However, the large number of GFP-positive cells precludes the identification of additional CX neurons and their individual trajectories.

### G10011-GFP positive neurons establish the FB primordium

CX neurons of holometabolus insects are born during embryonic and larval stages while much of the CX connectivity is established in the pupa. We examined the appearance of G10011-GFP labelled cells and the establishment of early CX connectivity at embryonic, larval and pupal stages. Firstly, we addressed G10011-GFP expression during embryogenesis. We observe the embryo staging nomenclature as suggested by Stollewerk which distinguishes 15 stages of neurogenesis: NS1(0%) to NS15 (100% neurogenesis) (Biffar and Stollewerk, 2014) for details refer to Figure **S2**). (Embryonic are given oordinations according to the body axis).

The earliest expression of G10011-GFP occurs at NS11 (65% of embryogenesis) in two small clusters of cells in anterior-medial positions of the brain (Figure 3A). Subsequently, cell numbers within these clusters increase and additional clusters form adjacently in more lateral positions (NS12, Figure 3B). In addition, small clusters GFP-positive cells appear in posterior-medial regions. Post-NS12, no significant increase of GFP-positive cells in the anterior-medial region takes place. By contrast, in posterior-medial and posterior-lateral regions multiple new GFP-positive cell clusters arise and early-born clusters gain in cell numbers (Figure 3C-D’, E,F and G,H). The strongest increase in GFP-positive cells is observed during stage NS15. At the end of embryogenesis each brain lobe contains an average number of 362 GFP-positive cells (n=4; Figure 2J), the vast majority of which reside in the medial area of the dorsoposterior brain. The stem cells of the brain generate their neural progeny in a stereotyped orientation towards the inside of the brain. Sequentially generated neurons remain close together such that lineage-related neurons appear like pearls on a string. GFP RNA in situ in stage NS15 G10011 embryos allows us to identify 14-16 “strings” of GFP-positive cells (Figure 6B’’). Taking into consideration that each embryonic brain hemisphere contains around 100 NBs (Biffar and Stollewerk, 2014), G10011-GFP is expressed in the progeny of about 15% of all NB lineages. Embryonic expression of G10011 outside of the brain is restricted to the stomodeum and the hindgut and GFP-expressing cell bodies are absent from the VNC (Figure 3K).

**Figure 3.**
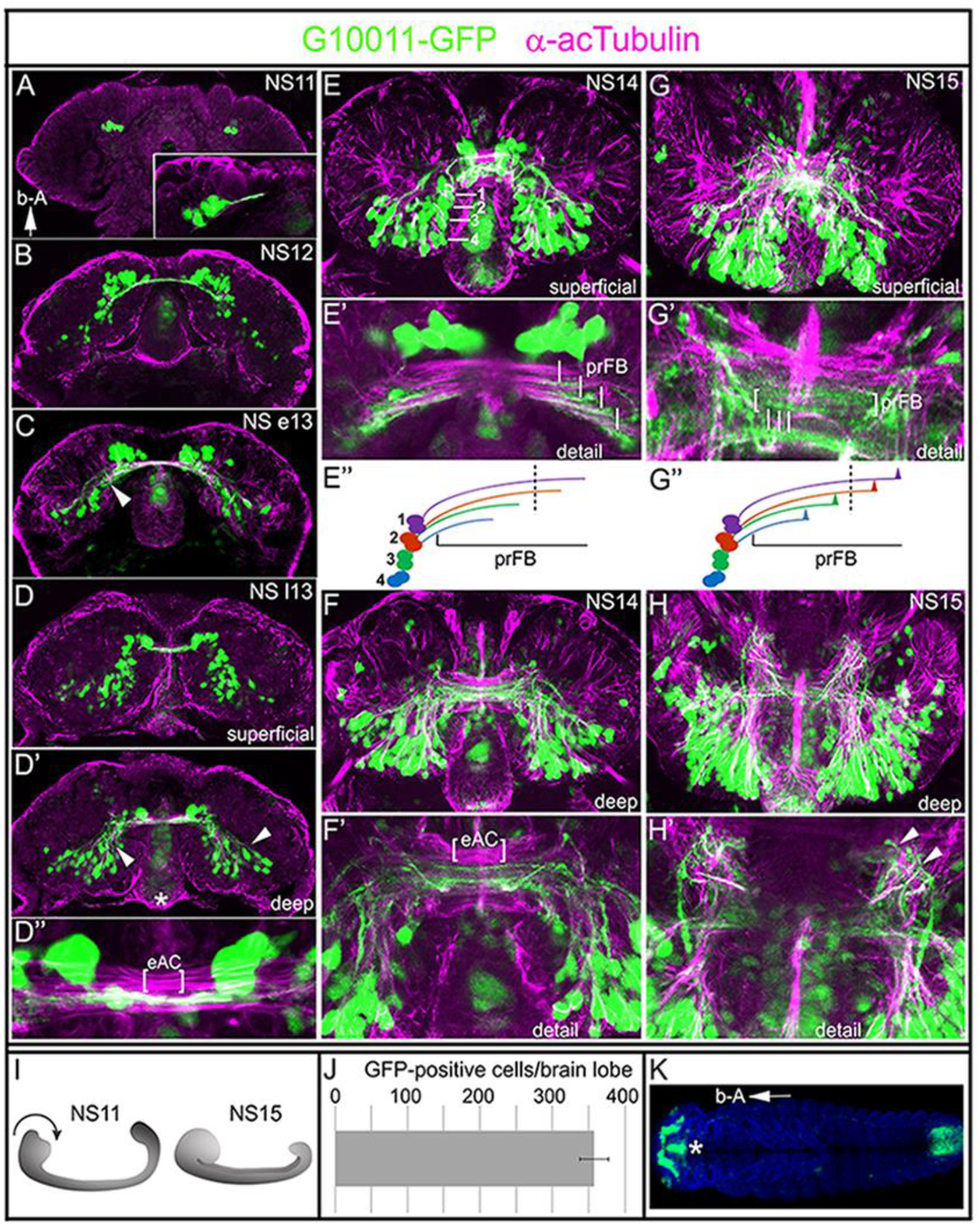
Embryonic expression of G10011 and formation of the embryonic commissural system. (**A-H’**) Developmental series of G10011-GFP brains beginning from stage NS11 (~65% embryogenesis) up to stage NS15 (100% embryogenesis). (For details of embryogenesis and staging see Figure **S2**) Coordinates are given according to the body axes (b-A arrow in A indicates ”anterior up”). (**A-H**) Double immuno-staining with α-GFP (green) and α-acetylated Tubulin (magenta) antibodies. (**A-C**) depict single confocal planes; anterior is up with respect to the body axis (arrow “b-A”). (**A, inset**) The first continuous commissural fascicle which links both hemispheres of the protocerebrum is established at late NS11. This primary commissural fascicle is strongly labelled by α-GFP. (**C**) Note that multiple GFP-positive cell clusters project their axons from posterior dorsomedial regions towards the primary brain commissure (arrow). (**D-D’**) Serial confocal sections were combined and visualized as maximum intensity projections to depict either superficial (**D**) or deep lying (**D’**) regions of a late NS13 brain. (**D’**) Multiple GFP-positive cell clusters project axons from posterior dorsomedial and dorsolateral regions towards the primary brain commissure (arrows). GFP also labels the stomodeum (asterisk). (**D’’**) Close-up of (**D’**) Multiple commissural fascicles have formed; only a subset is GFP-positive (eAC: embryonic anterior commissure). (**E-H’**) Serial confocal sections were combined and visualized as maximum intensity projections to depict superficial (**E-E’**) or deep (**F-F’**) lying regions of a stage NS14 brain. Note that nearly all GFP-positive cell bodies are located in the dorsoposterior brain. (**E**) White lines indicate four clusters of cells. We interpret these cells as the progeny of DM1-DM4 which differentiate into the columnar neurons. (**E’**) the four clusters of neurons produce four parallel running GFP-positive fascicles enter the commissural fiber system (white lines). We interpret these fibers as the precursors of the w, x, y, z tracts and hence as the prFB. (**E’’)** Schematic illustration: the trajectories of 4 cell clusters generate the prFB; dashed line indicates the ventral midline. (**F, F’**) GFP-positive input into the primary commissure stems largely from cells located in posterior dorsomedial and dorsolateral regions of the brain (arrowhead). (**G,G’)** superficial and (**H,H’**) deep lying regions of the NS15 brain. (**G**) Note the beginning defasciculation of GFP-positive commissural fiber tracts (white lines). (**G’’**) Schematic representation of the beginning defasciculation. (**H,H’**) Multiple GFP-positive fibers exist the brain and project towards the VNC (arrowheads). (**I**) Schematic representation of morphogenetic head movements during embryogenesis. (**J**) The average number of G10011-GFP-positive cells in late stage NS15 brain lobes is 362 (n=4). **(K**) Dorsal view of a whole-mount NS14 animal. Note that embryonic G10011-GFP expression is restricted to the brain, the stomodeum (asterisk) and the hindgut. Blue stain is DAPI. “b-A” arrow: anterior is left.

In the adult G10011 brain the columnar neurons of the FB are heavily labeled by GFP (Figure 2K). We asked whether these cells are of embryonic origin and establish the FB primordium (prFB) of the embryonic *Tribolium* brain. The prFB is formed by four contralaterally projecting fiber tracts which emanate from each brain hemisphere and constitute a part of the early commissural system (schematic illustration in Figure 3E’’). These fiber tracts are produced by four distinct neuronal neuroblasts (DM1-DM4) located in the posterior-medial brain (Andrade et al., 2019; Farnworth et al., 2020). To visualize the development of the commissural system, we co-stained G10011 embryos with *a*-acetylated Tubulin (acTub), currently the only available marker for *Tribolium* cell membranes. At late stage NS11 the first acTub-positive fascicle extends towards the midline. This fascicle is also labelled by G10011-GFP (Figure 3A, inset). From NS12 onwards, GFP-positive fiber tracts make numerous contributions to the commissural system (Figure 3B-D’’ and E-F). At stage NS14 GFP-positive fiber tracts form that are indicative of the prFB: four contralaterally projecting fiber tracts enter the commissural system as parallel tracts and fuse with the corresponding tracts of the opposing brain hemisphere (Figure 3E’,F). At late stage NS15 these fibers show the characteristic pattern of defasciculation which initiates the development of the columnar architecture of the FB (Figure 3G’, schematic illustration: G’’). In *Drosophila* it has been shown that the contralaterally projecting fibers which constitute the prFB pass through a channel formed by glial membranes (Andrade et al., 2019). We observed a similar arrangement in the *Tribolium* embryonic brain (Figure **S**3 A-A’’). Recently, we have shown that a subset of embryonically-born columnar neurons express the Retinal homeobox protein (Rx) (Farnworth et al., 2019). Double-staining with anti-GFP and anti-Rx revealed a partial overlap of Rx- and G10011-GFP expressing neurons (Figure **S**3 B-B’’’). Taken together, we conclude that G10011-GFP labels embryonically-born columnar neurons which establish the prFB.

In addition to columnar neurons, we identified subsets of AOTU- and ring neurons in the G10011-GFP adult brain that contribute to the CX. Studies in *Drosophila* have shown that some of these neurons are born in the embryo and persist to adulthood (Lovick et al., 2017; Bridi et al., 2019). Due to lack of specific markers we are unable to identify these cells unambiguously in the *Tribolium* embryo. However, in the late NS15 brain we observed GFP-positive cells which –based on cell number, morphology and location - we interpret as the AOTU cells (arrow) Figure **S3** C), as well as the putative “huging” expressing cells (arrowhead) and the neurosecretory cells of the prospective PI (arrow) (Figure **S3** D).

The adult VNC contains numerous GFP-positive longitudinal connectives which originate in the brain (Figure 1D; **S**1). We observed that longitudinal connectives arise during embryonic stages: several dorsomedially located cell clusters project axon tracts towards the VNC (Figure 3 H,H’). For a more detailed display of the major axon tracts in the late NS15 brain refer to Figure **S4**.

### G10011-GFP labels immature CX neuropils in the late Tribolium larva

During larval development, the brain strongly increases in size and undergoes major morphogenetic movements which together make it impossible to trace all embryonically-born G10011-GFP positive neurons to late larval stages. The number and distribution of GFP-positive cell bodies in the late larval brain (80-90% larval development) much resembles that of the adult brain (Figure 4A-L).

**Figure 4.**
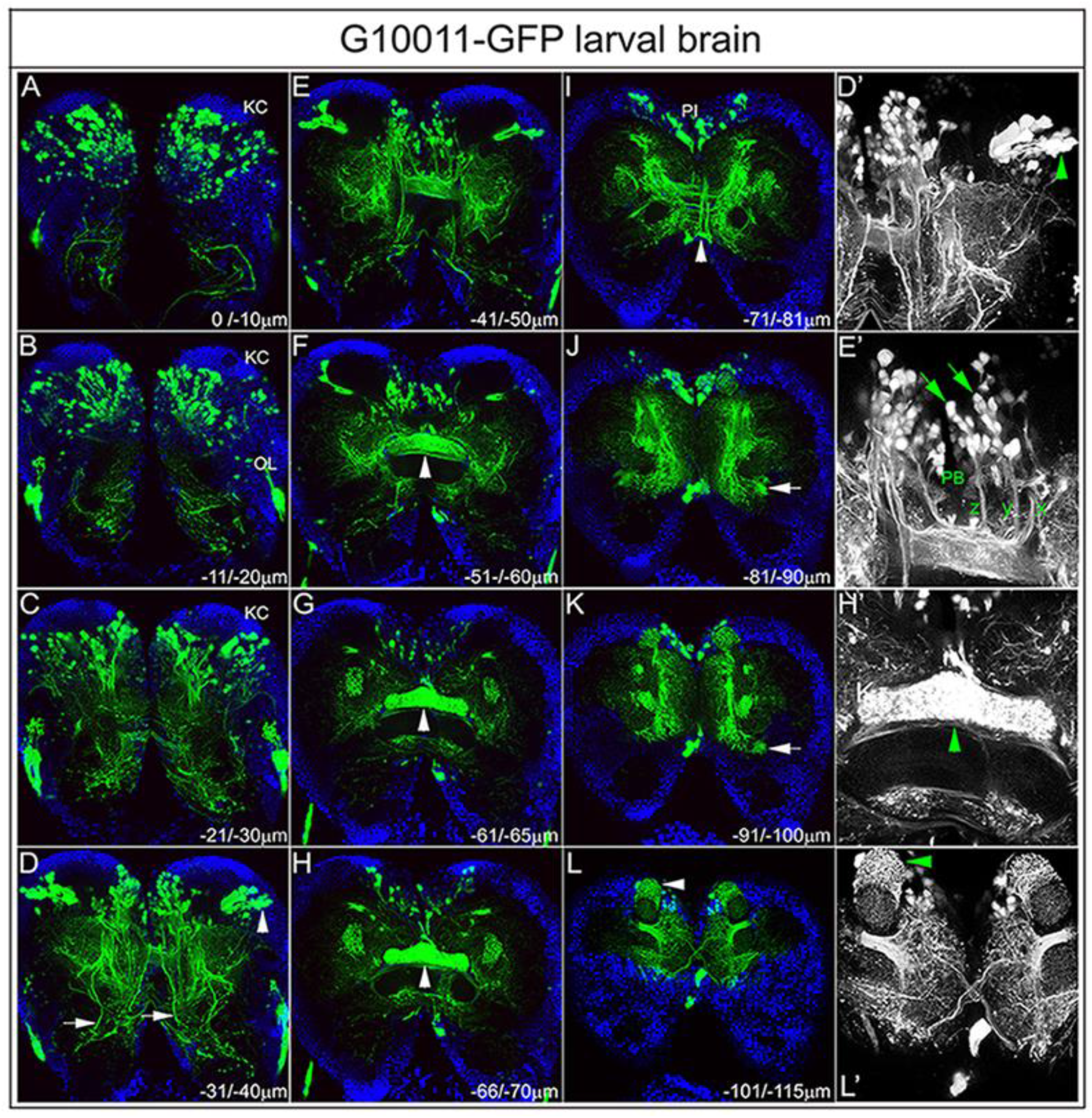
GFP expression in the late G10011 larva. (**A-L**) GFP auto-fluorescence (green) and DAPI (blue) staining. (**D’, E’, H’** and **L’**) GFP only. Serial confocal sections were combined and visualized as maximum intensity projections to display individual anatomical features. Scan direction is from the n-dorsal (**A**) towards the n-ventral (**L**) surface of the brain. Depth along the Z-axis is given in μm. (**A-C**) GFP-positive cell bodies in the posterior brain. Note that the Kenyon cells (KC) of the MBs do not express GFP. GFP fluorescence within the optic lobes (OL) may reflect the expression of the transfection marker 3xP3-GFP. (**D,D’**) Multiple axon tracts originating in the n-antero-medial and – lateral protocerebrum descend towards the VNC (arrows). Arrowheads indicate the dorsal and ventral clusters of AOTU neurons. (**E, E’**) A subset of columnar neurons (arrows) with their arborizations within the PB and their characteristic z, y, and x axon tracts (the w tract is not in focus). **(F-H, H’**) The FB is strongly labelled by GFP (arrowheads). Note that distinct elements of the EB are not yet present. (**I**) Ascending axon tracts originating from the putative “hugin-expressing” cells (arrowhead) project towards the PI. (**J-K**) GFP-positive ring neurons (arrows). (**L,L’**) GFP-expressing cells form multiple dendritic arborizations which enwrap distinct parts of the MBs (arrowheads).GFP-positive neurons reside nearly exclusively in the dorsoanterior brain. Most GFP-positive cells are located in the medial brain with exception of two large cell clusters which laterally abut the Kenyon cells (Figure 4D, D’). As in the adult, G10011-GFP expression is absent from the Kenyon cells of the MBs. Location, morphology and, in part, axon trajectories allow us to recognize sets of cells which we can also identify in the adult brain: notably, columnar neurons and a subset of AOTU neurons (Figure 4D-F,D’,E’), putative neurosecretory neurons of the PI (I,J), a subset of ring neurons (J,K) and the putative “hugin expressing” cells (K,L,L’).

In the late larva, the PB and the FB are clearly labeled by GFP. Fiber tracts emanating from the columnar neurons pass through individual glomeruli of the PB and then extend more ventrally to form characteristic tracts with multiple interhemispheric chiasmata before they extend further ventrally to build an immature FB within which a columnar structure is not yet obvious (Figure 4E’). The overall shape of the FB already resembles that of the adult FB but GFP fluorescence shows no obvious dorsoventral stratification at this stage (Figure 4G,H,H’). In the late larva, the EB is not yet detectable with G10011-GFP. In the late larval brain each hemisphere contains 384 GFP-positive cells (n=4). From larval stages onwards, we observe GFP fluorescence in several non-neural tissues (data not shown). Since these tissues are not easily accessible by RNA in situ hybridization, we currently do not know whether fluorescence reflects the bona fide expression of *TC007335* (see below) or is due to cryptic regulatory elements within our plasmid.

### G10011-GFP labels the PB, FB and EB in the late *Tribolium* pupa

In the late pupal brain the number and distribution of GFP-positive cell bodies is essentially the same as in the embryonic, larval and the adult brain (Figure 5A-C). The overall architecture of the pupal CX neuropils closely resembles that of their adult counterparts. The glomeruli of the PB are pronounced but fusion at the midline has not taken place as yet (Figure 5B). Within the FB, the columnar structure is well established (Figure 5A,B). In marked contrast to the late larval brain is the appearance of the EB with its characteristic radial segmentation (Figure 5C).

**Figure 5.**
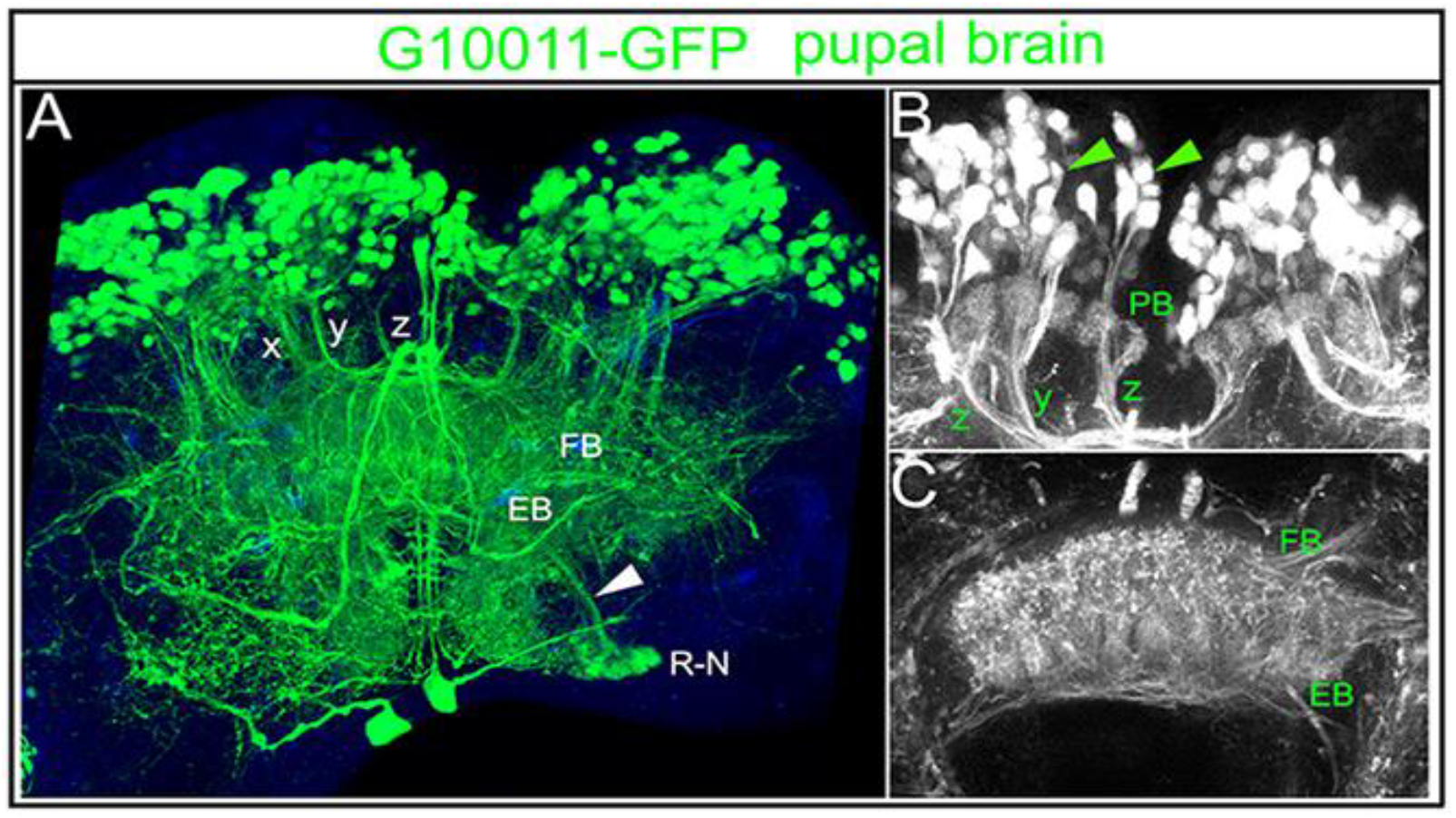
GFP-expression in the late (90% development) G10011 pupal brain. (**A-C**) GFP autofluorescence (in **A** combined with combined with DAPI staining, blue). (**A**) Confocal stack is visualized as maximum intensity projection. The columnar organization of the FB is well established. Note the ring neurons (R-N) and their projection towards the EB (arrowhead). (**B**) Columnar neurons, their arborizations within the glomeruli of the PB and their axon trajectories z, y, x (the w tract is not in focus). The PB is not yet fused at the midline. (**C**) The EB is well developed in the late pupa.

### G10011-GFP fluorescence reflects the RNA expression of the transcription factor TC-UNC-42

We mapped the plasmid insertion site of G10011 to the genomic position 6024777 within the first intron of *TC008169* (for mapping details see Figure **S**5). However, the expression of this gene did not match the one reported by G10011. Another candidate gene in this region is *TC007335* whose putative transcriptional start site is located 18.5kb upstream of the insertion site. To examine whether G10011-GFP reflects the expression of *TC007335* in the embryo, we performed fluorescent double-in-situ hybridization with a GFP and a *TC007335* RNA probe.

The GFP and *TC007335* signals colocalize at all embryonic stages, indicating that G10011-GFP faithful reports *TC007335* expression (Figure 6A-B’’ and data not shown). Furthermore, *TC007335* RNA in situ confirms that expression is restricted to the brain and stomodeum. We name *TC007335 shaking hands* (*skh*) to highlight the chiasma formed by cells of the PI (Figure 1B). *skh* encodes the ortholog of the *C.elegans* transcription factor UNC-42, a PRD-like homeodomain protein (Baran et al., 1999) (for a phylogenetic tree refer to Figure **S**6).

**Figure 6.**
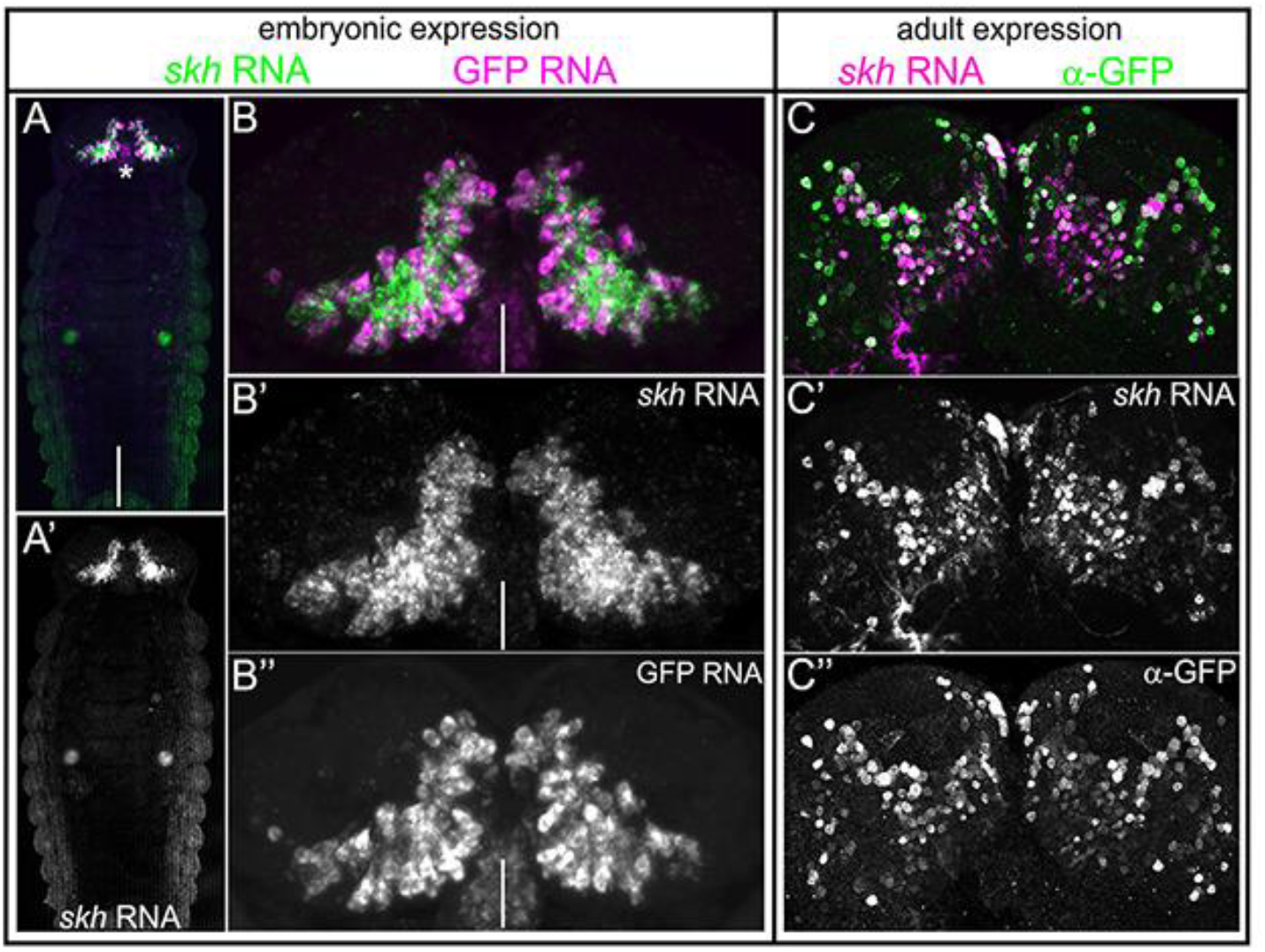
G10011-GFP reflects the RNA expression of Tc-*shaking hands (skh)* (TC007335). Double fluorescent in situ with a GFP (magenta) and a *skh* (green) RNA probe in a stage NS14 embryo. (**A-A’**) Dorsal view of a whole-mount embryo. Note that the expression of GFP and *skh* are restricted to the brain and stomodeum (asterisk). (**B-B’’**) GFP and *skh* RNA expression co-localize in the embryonic brain. White lines indicate the midline. (**C-C’’**) *skh* RNA in situ (magenta) combined with α-GFP antibody staining (green) in an adult G10011 brain. Serial confocal sections were combined and visualized as maximum intensity projections. Note the co-localization of *skh* RNA and GFP protein.

G10011-GFP fluorescence is still strong in the adult brain. To investigate whether this reflects GFP perdurance or the continued expression of *Tc*-*skh*, we performed *skh* whole-mount *RNA* in situ combined with α-GFP staining in the adult brain. All GFP-positive cells are also *skh RNA* positive, demonstrating the continued expression *of skh* (Figure 6C-C’’).

### *Tc-skh* expression is restricted to neurons

Embryonic and larval brains contain mitotically active and postmitotic cells. Mitotically active cells are NBs and their immediate progeny. To determine whether Tc-*skh* is expressed in mitotically active cells, we double-stained embryos and larval brains with α-GFP and the mitosis marker α-Phospho-histone-3 (PH-3).

Co-localization of α-GFP and α-PH-3 signals is never observed, indicating that *Tc*-*skh* expression is restricted to postmitotic cells (Figure 7 A-A’’ and data not shown). This conclusion is supported by the observation that GFP-fluorescence is absent from the superficial, neuroblast, layer of the brain. To examine whether *Tc*-*skh* is expressed in glia, we double-stained embryos and adult brains with α-GFP and the glial marker α-Repo. α-GFP and α-Repo signals do not co-localize (Figure 7B-C’’ and data not shown). We conclude that *Tc*-*skh* expression is restricted to neurons.

**Figure 7.**
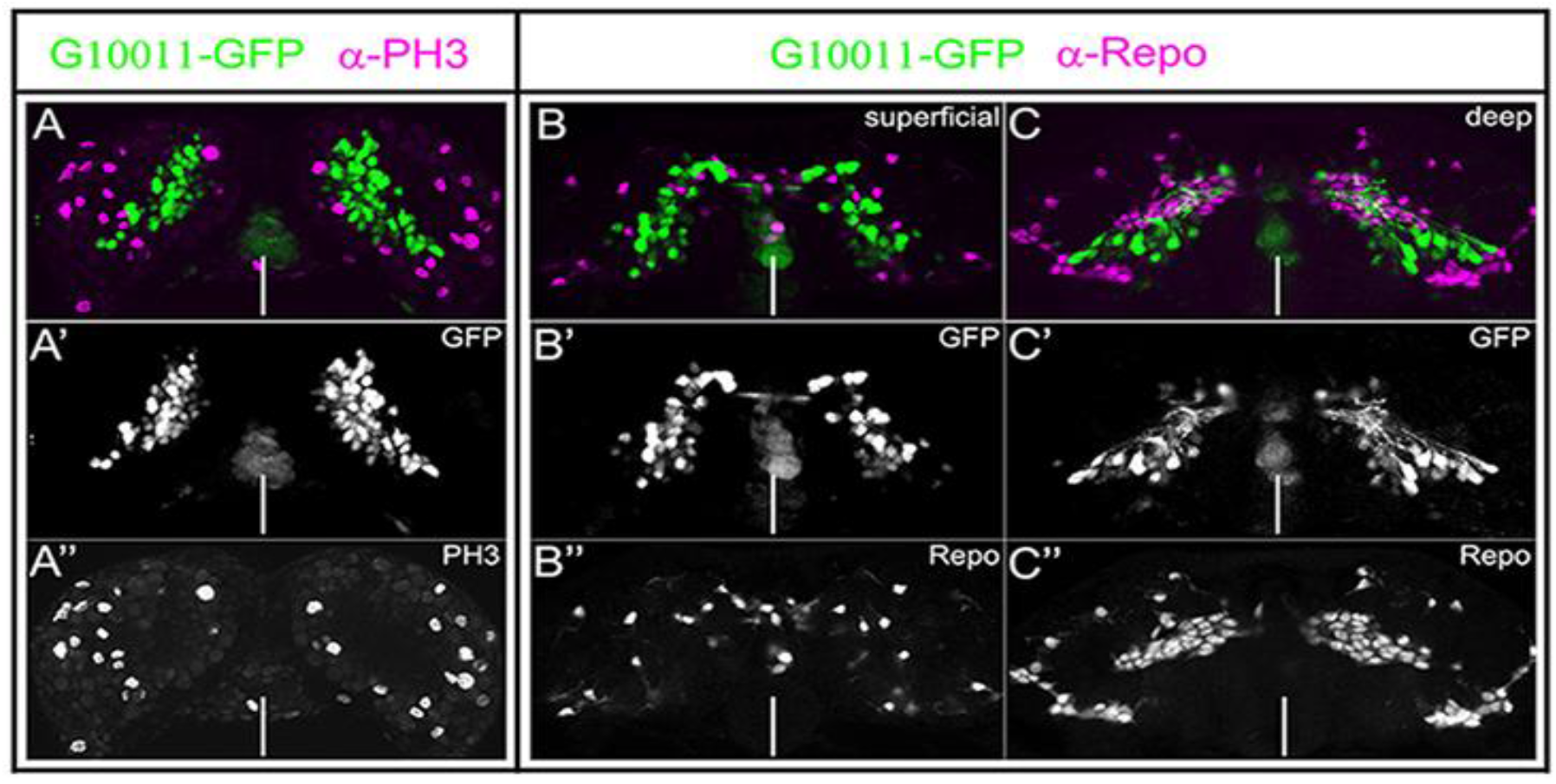
G10011-GFP expression is restricted to non-dividing, non-glial cells. Embryonic G10011 brains were stained with α-GFP (green) and α-PH3 (**A-A’’**) or α-Repo (**B-C’’**) (magenta). Serial confocal sections were combined and visualized as maximum intensity projections. (**A-A’’**) NS13 and (**B-C’’**) NS14. Note that there is no overlap of GFP- and PH3- or Repo-expressing cells. White lines indicate the midline.

### *Tc-skh* knock-down results in axon outgrowth defects and a reduction of GFP-positive cells

To explore the effects of reduced Tc*-*Skh function in the embryo, we performed parental RNAi in G10011 animals using two non-overlapping dsRNA fragments (“frag1” and “frag2”). Knock-down phenotypes were examined by double staining with α-GFP and α-acTubulin.

Loss of Tc-Skh has drastic consequences for the outgrowth of all G10011-GFP positive axons: in severely affected embryos, no contra-laterally projecting axons enter the commissural system and hence, no prFB is formed (Figure 8A-D’’). Axon outgrowth defects occur with high penetrance: RNAi with “frag1” and “frag2” results in severe defects in 71% and 48% of the embryos, respectively. Examination of GFP-fluorescent cells shows that some axon outgrowth still takes place but axons terminate prematurely close to the respective cell bodies (Figure 7C). Axon extension defects are restricted to GFP-positive trajectories: acTubulin positive but GFP negative axon trajectories form normally (Figure 7C,D’’). We conclude that the requirement of Tc-Skh for axon extension is cell-autonomous.

**Figure 8.**
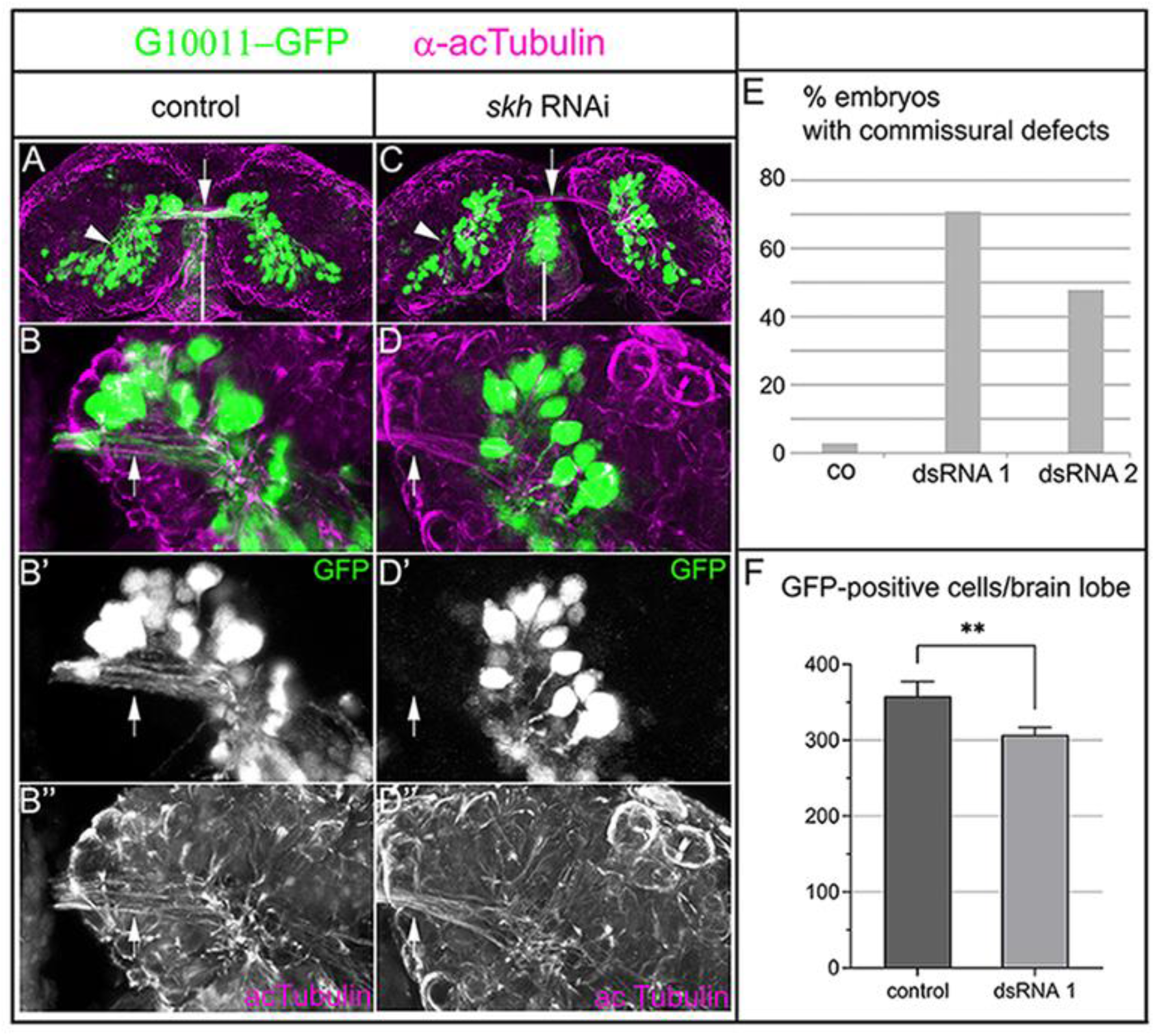
parental RNAi of *skh* leads to severe axon outgrowth defects in the embryonic brain. G10011-GFP stage NS14 brains double-stained with α-GFP (green) and α-acTubulin (magenta); dorsal views. (**A-B’’**) Control brain (progeny of buffer-injected pupae). (**A**) whole brain at low magnification; GFP-positive axon tracts join the commissural system linking both hemispheres of the protocerebrum (arrow). The arrowhead indicates GFP-positive cell clusters in the posterior brain. (**B-B’’**) close-ups; GFP-positive axons project towards the midline (arrows). GFP-positive input into the primary commissure stems largely from cells located in posterior dorsomedial and dorsolateral regions of the brain (arrow). (**C-D’’**) *skh* RNAi brain. (**C**) whole brain at low magnification; GFP-positive axons fail to join the commissural system. Arrowhead indicates the loss of GFP-positive cell clusters. (**D-D’’**) Close-ups; (**D’**) GFP-positive axons stall while most acTubulin-positive axons are unaffected (**D’’**). White lines indicate the midline. (**E,F**) Quantification of *skh* RNAi phenotypes. (**E**) commissural defects were scored at stages NS14 and NS15. Buffer-injected control (co) n=80 (2 biological replicates) 3% defects, dsRNA frag1 n=85 (2 biological replicates) 71% defects, dsRNA frag2 n=35, 48% defects. (**F**) Loss of GFP-positive cells in *skh* RNAi embryos were scored at NS15. Buffer-injected control (co): 362 GFP-positive cells (n=4), dsRNA frag1: 308 GFP-positive cells (n=4). Statistical significance was determined by one way ANOVA, ** = P<0.05.

In addition to axonal defects, we observe a moderate reduction of GFP-positive cells in knock-down embryos (compare Figure 8A with C; quantification in 8F). Due to the lack of specific markers for G10011-GFP positive cells, we are unable to determine whether loss of Skh is due to apoptosis or reflects an auto-regulatory feedback loop in the maintenance of Skh expression.

### The embryonic expression patterns of *Tribolium* and *Drosophila-skh* are conserved

The *Drosophila* ortholog of *Tribolium skh* is encoded by CG32532 (see Figure **S**6). Its gene product is one of the few homeodomain proteins which have remained uncharacterized. We examined the embryonic expression pattern of *Dm*-*skh* by RNA in situ hybridization (Figure 9A-E’).

**Figure 9.**
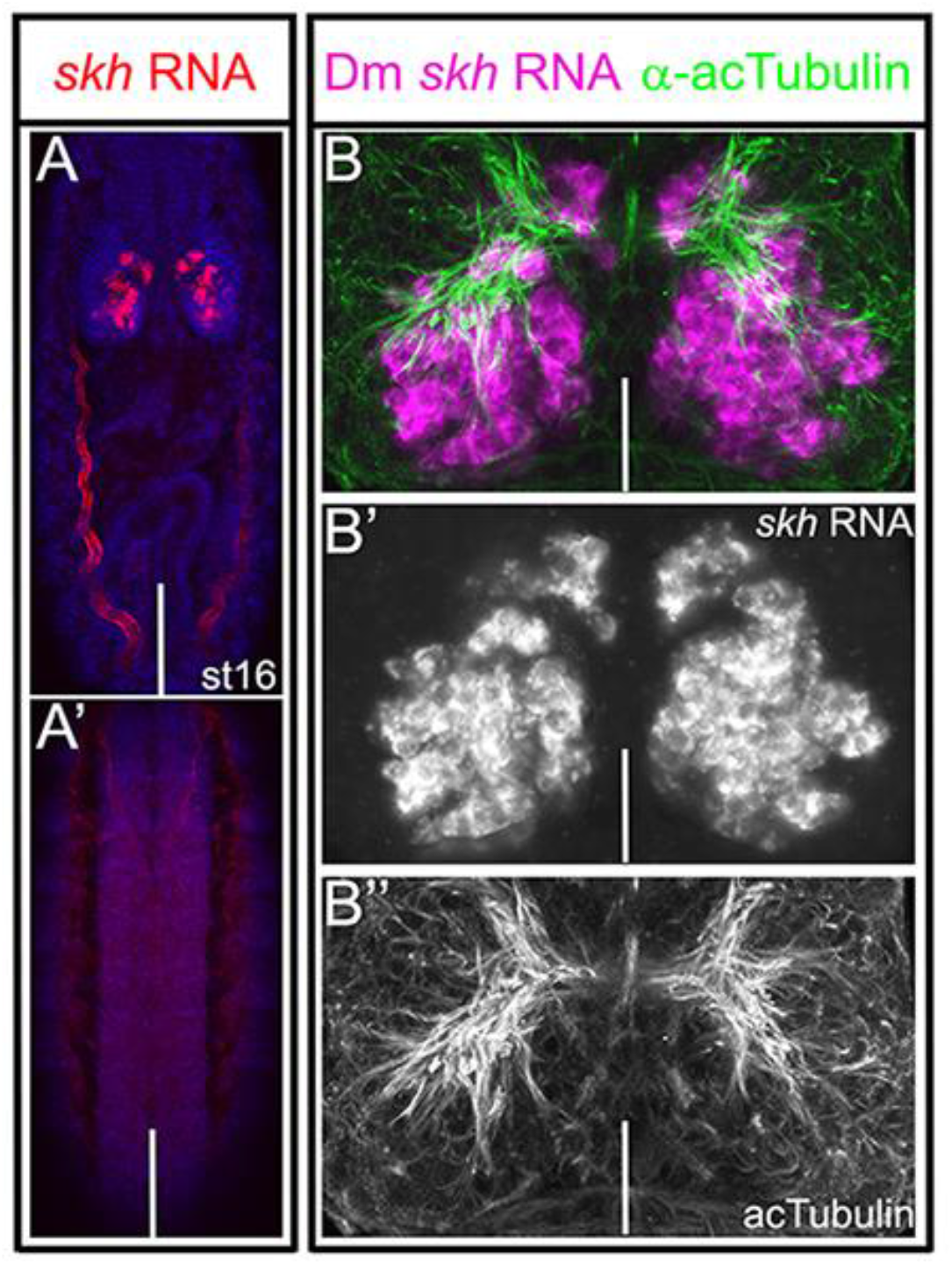
Embryonic expression of Dm *skh*. (**A,A’**) *skh* RNA in situ (red) and DAPI staining (blue). (**A,A’**) stage 16 whole-mount embryo; anterior is up. (**A**) dorsal view; (**A’**) ventral view. Note that Dm *skh* RNA is restricted to the brain. Red fluorescence in the trachea is an in situ artefact (arrow). (**B**) Spatial organization of *skh* RNA expressing cells and major axon tracts; *skh* RNA in situ (magenta) combined with an α-acetylated Tubulin (green) staining. (**B’)***skh* RNA only. Compare with Figure 6B: embryonic *skh*-expressing cells are similarly distributed in *Tribolium* and *Drosophila*. (**B’’**) α-acetylated Tubulin only. White lines indicate the dorsal midline in all panels except (**A’**) where it marks the ventral midline.

As its *Tribolium* ortholog, *Dm-skh* is expressed in the brain but is absent from the VNC and non-neural tissues (Figure 9A,A’).To compare the spatial arrangement of *Drosophila* and *Tribolium skh* positive cells, we double-labeled *Drosophila* embryos with *skh* RNA and α– acTubulin and examined the positions of cell bodies relative to the commissural system. At the end of embryogenesis, the spatial arrangement appears highly similar in both organisms (compare Figure 9B-B’’ with Figure 6B’): small clusters of *skh* positive cells are located just anteriorly to the commissural system while the vast majority of cells reside posteriorly to the commissural system in the dorsomedial brain. In a few cases we are able to follow the trajectories of *skh* positive cells and find that some of them enter the commissural system (Figure 9B-B’’).

## Discussion

### G10011-GFP is a useful tool for the study of the dynamics of CX development

Although the anatomy of the adult CX is well described in many insect species, the CX is vastly understudied from a comparative developmental perspective. Beside a large body of work addressing CX development in *Drosophila*, only the grasshopper has been thoroughly investigated as an alternative insect model (Boyan and Williams, 2011; Boyan and Reichert, 2011; Boyan et al., 2017). Developmental studies in non-*Drosophila* models are hampered by a lack of anatomical information at the single cell level but also by a near complete lack of molecular and genetic tools. We seek to establish *Tribolium* as an alternative insect model to study CX development (He et al., 2019; Farnworth et al., 2020). The line G10011-GFP labels several neuropils of the adult CX but does not label other major neuropils, e.g. the mushroom bodies and the antennal lobes. Our results suggest that many GFP-positive CX neurons are born early in development, making G10011-GFP a useful tool for the study of the dynamics of CX formation: indeed, G10011-GFP expression confirms and extends earlier findings that the *Tribolium* FB is largely assembled in the larva, while a distinct EB forms later in the pupa (Panov, 1959; Koniszewski et al., 2016; Farnworth et al., 2020). In combination with additional markers, G10011-GFP will be a valuable tool to identify and characterize a subset of CX neurons at the single cell level, thereby contributing to a much needed *Tribolium* brain atlas.

The *Drosophila* ortholog of Tc-*skh* shows an RNA pattern in the embryo that is highly similar to that of *Tc*-*skh* suggesting that early expression is conserved. The generation of a corresponding imaging line should provide a means for a comparative study of *Drosophila* and *Triboliu*m CX development at the anatomical and the molecular level.

### Tc-Skh is a putative terminal selector of neuronal subtype identity

Terminal selector expression in neurons is continuous from cell birth to cell death. Therefore, such factors provide excellent markers for specific subsets of neurons for developmental, molecular and evolutionary studies. With this work, we identify Skh as the first putative terminal selector in neurons that contribute to the CX. Tc-Skh is the ortholog of *C. elegans* Unc-42 whose role in the specification of neuronal subtypes is well described (Wightman et al., 1997). *unc-42* (*<u>unc</u>oordinated - 42*) was first discovered by Brenner in his classic screen of mutants which show abnormal locomotion (Brenner, 1974). A later study showed that Unc-42 is required for axon pathfinding in a subset of neurons which facilitate a specific locomotor routine (Baran et al., 1999). Studies of *C. elegans* Unc-42 and other transcription factors have led to the concept of terminal selectors as regulators of neuronal subtype identity (Hobert, 2008). In contrast to developmental genes that are expressed early in the gene regulatory cascade, terminal selectors are the final targets of the cascade. Maintenance of terminal selector expression is accomplished by positive auto-regulatory feed-back loops; accordingly, loss of terminal selector activity results in the loss of terminal selector gene expression at later stages. The lifelong expression of terminal selectors facilitates the regulation of early aspects of subtype differentiation, such as axon pathfinding, as well as late aspects like the maintenance of structural and molecular features of the mature neuron.

Our study shows that *Tribolium skh* expression is characteristic for terminal selector genes. *Tc*-*skh* is not expressed in progenitor cells but is restricted to postmitotic neurons. We hypothesize that the adult expression of *T*c-*skh* reflects the lifelong expression in many embryonically-born neurons; however, due to the lack of genetic tools for permanent cell marking, we can demonstrate this only for embryonically-born columnar neurons and neurons of the PI which can be traced to adulthood. An early aspect of columnar neuron identity is their axonal projection which leads to the establishment of the prFB. Knock-down of Tc-Skh abolishes columnar (and other) neuron axon extensions indicating that a requirement of Skh for the development of proper connectivity is conserved between *Tribolium* and *C. elegans*. In late *Tribolium* knock-down embryos we observe a moderate loss of G10011-GFP fluorescence suggesting that maintenance of *Tc-skh* expression by an auto-regulatory feed-back loop may be another conserved feature.

The term terminal selector derives from studies in *C. elegans* where certain transcription factors directly co-regulate differentiation genes which together bring about all the specific features of a distinct neuronal subtype. The target genes of *Tc*-*skh* are currently unknown. In early development they are likely to include differentiation genes required for axon outgrowth/pathfinding like cell adhesion molecules and receptors for guidance cues. The question of whether *Tc*-*skh* coordinately directs the expression of a battery of effector genes at any stage during the life of a neuron remains to be investigated.

*Tc*-*skh* expression is not restricted to one particular neuronal subtype but found in many neurons with different morphological features. Therefore, we expect additional transcription factors to act in parallel to or in combination with Tc-Skh to specify distinct identities. Combinatorial action in the regulation of effector genes is a common theme in *Drosophila* neurons (Allan and Thor, 2015). Recently identified transcription factors that are expressed in embryonic *Tribolium* CX neurons, Tc-FoxQ2 and Tc-RX, are good candidates for factors acting in combination with Tc-Skh (He et al., 2019; Farnworth et al., 2020).

The activation of terminal selector gene expression is the final step of hierarchical gene regulatory cascades which provide early spatial and temporal information for the specification of subtypes. Our observation that most *Tc*-*skh* expressing cells arise in the posterior dorsomedial brain suggests that relevant spatial cues mark this region. A good candidate is SIX3/OPTIX: SIX3 is expressed in the neuroectoderm from which a large part of the dorsomedial brain derives. Moreover, knock-down of *six3* results in severe defects in the embryonic CX (Posnien et al., 2011). By contrast, a role for a temporal cascade in neuronal subtype specification in *Tribolium* has not been shown as yet.

*C. elegans unc-42* expression is not restricted to interneurons, called command neurons, but also occurs in sensory neurons which provide incoming information and motorneurons which facilitate the output. Together a subset of UNC-42-positive neurons forms a distinct circuit which facilitates a specific locomotor routine. At present, we do not know whether *Tc*-*skh* is expressed in sensory neurons which could provide incoming information to the CX. G10011-GFP fluorescence is absent from the VNC suggesting that *skh* is not expressed in motor neurons. A study of Tc-Skh expression in sensory and motor neurons will have to await the generation of a specific antibody.

In addition to *skh*, a number of other transcription factors have been shown to act as terminal selectors in *C. elegans* neuronal subtype specification (Hobert, 2016). Determining their expression patterns and their target genes in *Tribolium* and *Drosophia* CX neurons will contribute to a better understanding of CX formation and may uncover a molecular basis for anatomical differences of the CX in these species.

## Material and Methods

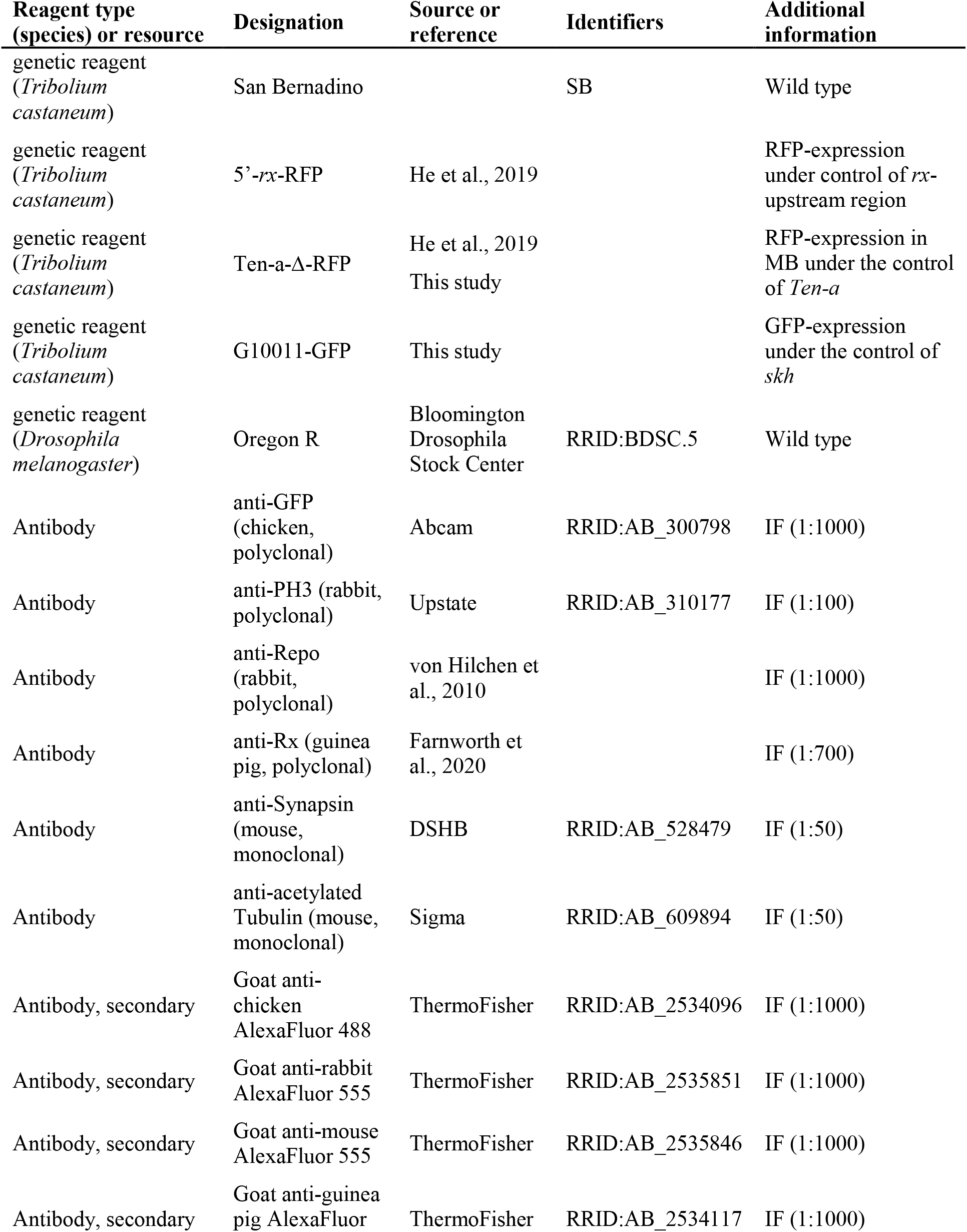
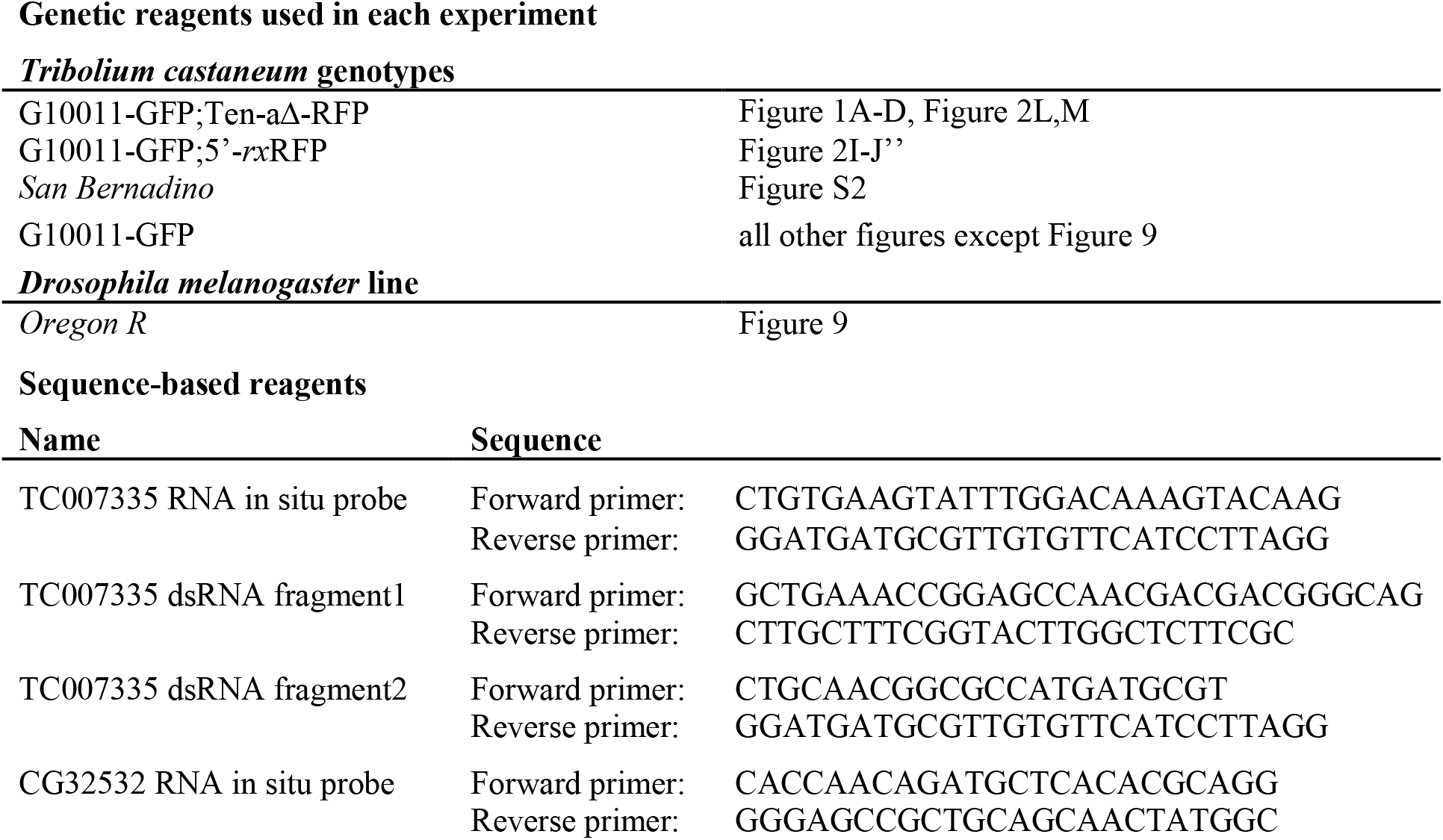
Key resources table.

### Animal husbandry

*Tribolium castaneum* (NCBI:txid7070) beetles were maintained on standard wholemeal wheat flour (type 1050) at 28°C. To obtain embryos, the beetles were transferred to fine wheat flour (type 405) and kept at 32°C. Egg lay was allowed for 24hrs. Subsequently, the embryos were separated from the beetles, aged for an additional 24-48hrs at 32°C and then collected for fixation. The *San Bernadino* strain was used as wild type.

*Drosophila melanogaster Oregon R* flies were maintained at 18°C on standard cornmeal agar supplemented with dry yeast flakes. To obtain embryos, flies were placed in collection cages with apple juice-agar plates smeared with fresh yeast paste and placed at 25°C. Egg lay was allowed for 4hrs. Then, the apple juice plates were removed from the cages, aged for an additional 16hrs at 25°C and then collected for fixation.

### Fixation

*Tribolium* embryos were collected, fixed and stored as described (Buescher et al., 2020). *Tribolium* larval, pupal and adult brains were dissected in ice-cold 1x phosphate-buffered saline (PBS) for up to 30 min. Then methanol-free formaldehyde was added to a final concentration of 4% (v/v). Fixation was performed for 30 (larval-), 45 (pupal-) or 60 (adult-brains) min on ice. Subsequently, the brains were washed 3x for 20 min each with ice-cold 1x PBST (PBS including 0.1% Triton-X100). In the second wash, DAPI was added to a final concentration of 1ng/μl. Brains not dedicated to immunohistochemistry were mounted in VectaShield H-100 and imaged immediately. Brains dedicated to immunohistochemistry were placed into blocking solution containing 3% (w/v) bovine serum albumin (BSA) and 0.05% sodium azide. Adult brains dedicated to RNA in situ hybridization were dehydrated by putting them through an ethanol series: 25% ETOH:75% PBS, 50% ETOH:50% PBS, 75% ETOH:25% PBS for 5 min incubation each. Finally, the brains were placed in 100% ETOH and kept at −20°C for several days prior to in situ hybridization.

### Immunohistochemistry

*Tribolium* and *Drosophila* embryos: Methanol was discarded from the fixed embryo collections. Subsequently, the embryos were washed 3x with 1x PBST for 20 min each at RT. Embryos were blocked for 1-2hrs in 3% BSA (w/v) (containing 0.05% sodium azide) at RT. Primary antibodies were added at the indicated concentrations and incubation was performed overnight on a rotating wheel at 4°C. Then the primary antibodies were removed and the embryos were washed 3x with 1x PBST for 30 min each at RT. Secondary antibodies were added at a dilution of 1:1000 and incubation was performed on a rotating wheel for 90 min at RT. Subsequently, the embryos were washed 3x for 20 min each with 1x PBST (PBS containing 0.1% Triton-X100). In the first wash, DAPI was added to a final concentration of 1ng/μl. Finally as much liquid as possible was removed and VectaShield was added.

*Tribolium* germ bands were freed of yolk with the help of a fine brush, mounted with the dorsal side up and imaged. *Drosophila* embryos were pipetted onto microscope slides and imaged as whole-mounts.

Immunohistochemistry with adult brains was performed essentially as with embryos except for the following modifications: the concentration of Triton X-100 was raised to 0.5%, the incubation period with the primary antibody was extended to about 40hrs and incubation with the secondary antibody was performed overnight at 4°C.

See Table 1 for a list of all primary and secondary antibodies used in this study.

### FISH

Single- and double-fluorescent RNA in situ hybridization and RNA in situ hybridization followed by antibody staining of *Tribolium/Drosophila* embryos was performed as described (Buescher et al., 2020). To generate a *skh* specific RNA in situ probe, a DNA fragment was generated by PCR using wildtype embryonic cDNA (*Tribolium/Drosophila*) as templates (for details see list of reagents). The PCR products were cleaned up by gel-electrophoresis, extracted and used as template for an additional round of amplification using the same gene-specific primer pairs but with the modification of an added T7 RNA transcriptase binding site at the 5’end of the reverse primer. Dioxigenin- or fluorescein-labeled RNA probes were produced using the Roche RNA labelling kit. For RNA in situ hybridizations, the probes were used at a concentration of 4ng/μl (total hybridization volume: 50-100 μl).

RNA in situ hybridization in *Tribolium* adult brains was performed essentially as in embryos, except for the following modifications: the concentration of Triton X-100 was raised to 0.5%, the RNA hybridization period was prolonged to 48hrs and incubation with the respective antibodies was performed for 48hrs at 4°C.

### Image acquisition

Confocal serial scanning images were acquired at 1.5-2μm intervals using a LSM 510 microscope (Carl Zeiss) using either a 20x 0.5 Plan-Neofluar or a 40x 1.4 Plan-Neofluar objective (Carl Zeiss). Stacks were processed using the Zeiss LSM Browser software and whole or parts of stacks were visualized as maximum intensity projections. Brightness, contrast, size and resolution of the images were processed in Adobe Photoshop CS. The video image of the G10011 adult brain was generated with the ImageJ software (Figure S1).

### Nomenclature used in anatomical analysis

For *Tribolium* and *Drosophila* embryos the axes used for anatomical analysis in this study are the body axes (in Figure 3A and K “b-A” indicates anterior with respect to the body axis).

For *Tribolium* postembryonic stages (larva, pupa, adult) the axes used for anatomical studies are neuraxes. According to the neuraxes, the protocerebral bridge and the fan-shaped body are located n-dorsal of the ellipsoid body. For a detailed description of the body and the neuraxes in *Tribolium* and *Drosophila* refer to Farnworth et al., 2020.

### Insertion site mapping of the enhancer trap line G10011

The genomic location of the plasmid insertion was determined by inverse PCR (Thibault et al., 2004). Genomic DNA was extracted from 3 beetles following a standard protocol. The genomic DNA was digested with the restriction enzyme Sau3A, highly diluted and ligated under conditions which facilitate an intramolecular circularization. Subsequently, the ligation products were amplified by PCR using plasmid-specific primers. The PCR product was cleaned up by gel-electrophoresis, extracted and sequenced. Blasting of the sequence against the *Tribolium* genome (genome release 3.0) indicated the insertion of the plasmid on chromosome 4 at the genomic position 6024777, which is 18.5 kb upstream of the predicted gene *TC007335* (transcription start site 6006266). Using double-fluorescent in situ hybridization, we confirmed that the GFP-expression of G10011 faithfully reflects the RNA expression of *TC007335* in the embryo and the adult brain. Sequence analysis of the predicted coding region indicates that *TC007335* encodes a paired-like homeodomain transcription factor and is the ortholog of C. elegans *unc-42* and *Drosophila CG32532*. We name *TC007335* and *CG32532 Tc*- and *Dm*-*shaking hands* (*skh*), respectively.

### *skh* knock-down

parental RNAi in *Tribolium*: 300-400 female G10011 pupae (at 70-80% pupal development) were injected with dsRNA (2μg /μl) or injection buffer only (control) using a FemtoJet Express (Eppendorf). Injected pupae were placed on fine wheat flour (type 405) for 24hrs at 28°C. Eclosed beetles were added to approx. 200 male G10011 beetles and maintained for another 24hrs at 28°C. Then all beetles were collected, placed on fresh fine wheat flour and shifted to 32°C. Eggs were collected for 24hrs, aged for an additional 48hrs and then fixated for immunostaining. Eggs were collected for 8 consecutive days. Pupal injections were performed twice with fragment 1 and once with fragment 2.

### Generation of gene-specific dsRNA fragments

Embryonic cDNA (0-72hrs), prepared from the *San Bernadino* wildtype strain, was used as template for the generation of gene-specific fragments within the predicted *TC007335* transcribed region. Two primer pairs were used to generate two non-overlapping fragments (fragment 1: 287bp, fragment 2: 261bp) by PCR (for details: see list of reagents). The products were cleaned up by gel-electrophoresis, extracted and used as templates for an additional round of amplification using the same gene-specific primer pairs but with the modification of added T7 RNA transcriptase binding sites at both 5’ends. The PCR products were used as templates for large scale RNA synthesis using the MEGAscript T7 Transcription Kit (Invitrogen). The dsRNA was precipitated with LiCl, washed with 70% ethanol, dried and dissolved in injection buffer (1.4mM NaCl, 0.07mM Na_2_HPO_4_ 0.03mM KH_2_PO_4_, 4mM KCl, pH6.8) to a concentration of 2μg/μl.

### Statistics

Data represent mean +/− standard deviation. Sample size in each experiment =4. In Figure 8, *skhRNAi*, the sample size n= 35-85. Figure 8F: statistical significance was determined with one-way-Anova. **=P<0.05.

## Acknowledgements

We thank Elke Kuester for excellent technical assistance, Dr. Gerd Vorbrueggen for his continued interest in the project and critical reading of the manuscript and Dr. Benjamin Altenhein for providing the α-Repo antibody. The monoclonal antibody α-SYNORF1 was obtained from the Developmental Studies Hybridoma Bank, created by the NICHD of the NIH and maintained at The University of Iowa, Department of Biology, Iowa City, IA 52242. We acknowledge support by the Open Access Publication Funds of the University of Goettingen.

## Contributions

Concept: MB. Investigation: MB, NGP. Methodology: MB. Formal analysis: MB, NGP. Validation: MB, NGP. Resources: MB, GB. Writing, original draft: MB. Writing, review and editing: MB, NGP, GB. Supervision: MB, GB. Funding: GB.

The authors declare that no competing interests exist.

