## supplemental figures for "Shaking hands is a putative terminal selector and controls axon outgrowth of central complex neurons in the insect model *Tribolium*"

**Figure S 1**

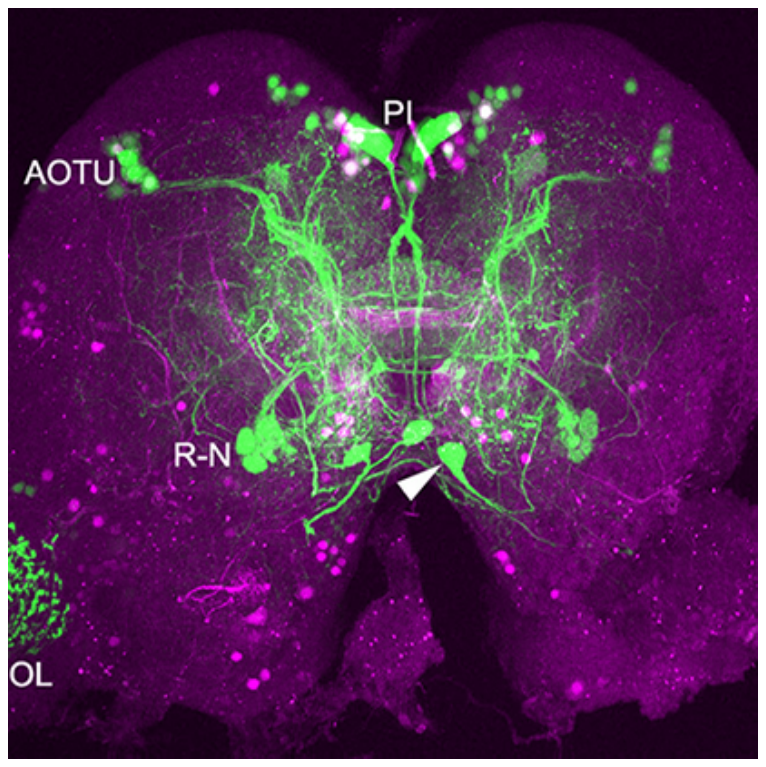

**Figure S 1:** G10011-GFP; 5’-rx-RFP adult brain. PI: pars intercerebralis, AOTU: neurons of the anterior optic tubercle, R-N: ring neurons, OL: optic lobe, arrowhead: putative huging expressing neuron. An animated 3-D version can be found at <https://figshare.com/projects/Additional_data_for_Garc_a_P_rez_et_al_Tribolium_shaking_hands_is_a_putative_terminal_selector_and_controls_axon_outgrowth_of_central_complex_neurons/93149>

**Figure S 2**

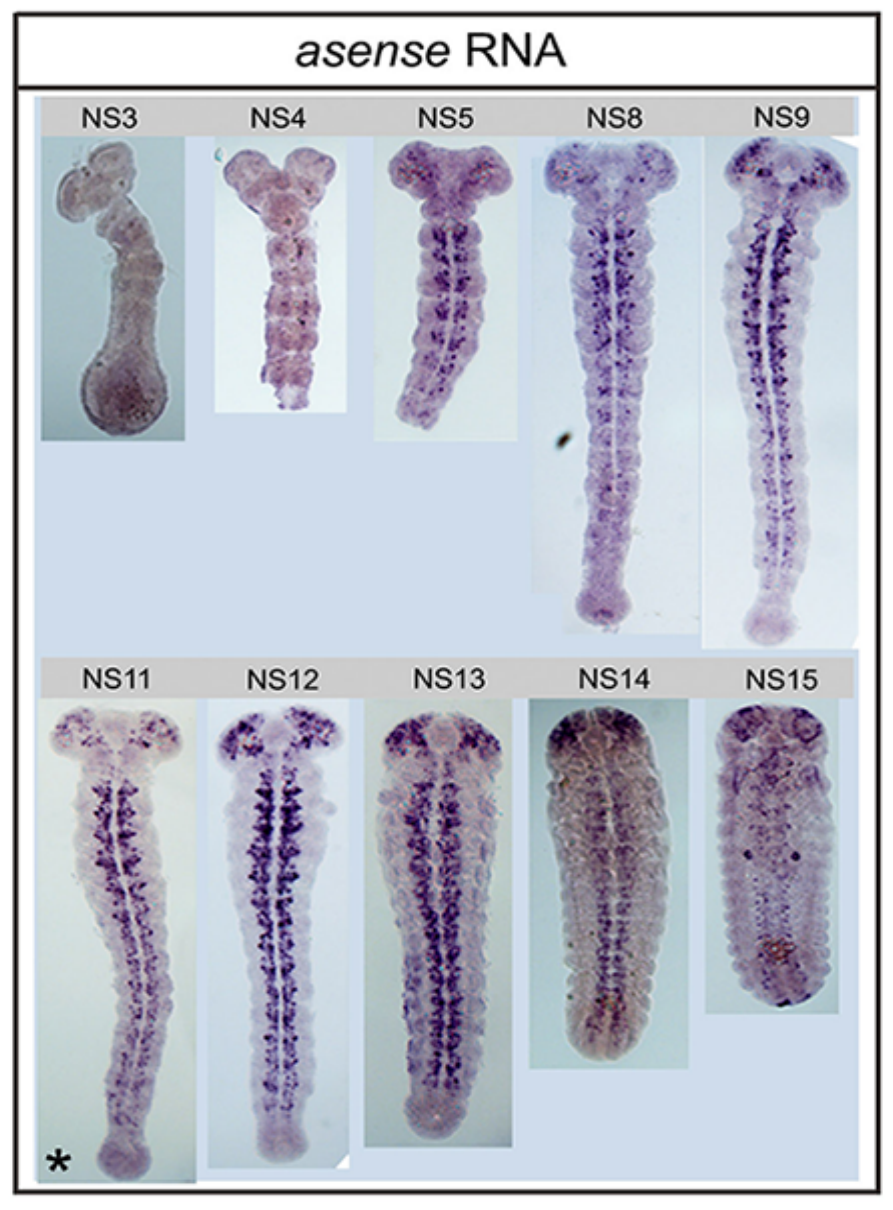

**Figure S 2:** Embryonic development of *Tribolium*; dorsal view of whole-mount embryos. Neuroblasts (NBs) are visualized by *asense* RNA in situ. *Tribolium* embryogenesis is subdivided into 15 stages from NS1 (0% embryogenesis) to NS15 (100% embryogenesis). NB formation in the brain begins at stage NS4. Subsequently, the number of NBs increases steadily until stage NS13/NS14. The precise number of brain NBs that are generated during embryogenesis is not known but is estimated to be around 100 per lobe. G100111-GFP-positive cells are first observed at stage NS11; that is at approx. 60% of embryonic development (stage11 is marked by an asterisk).

**Figure S 3**

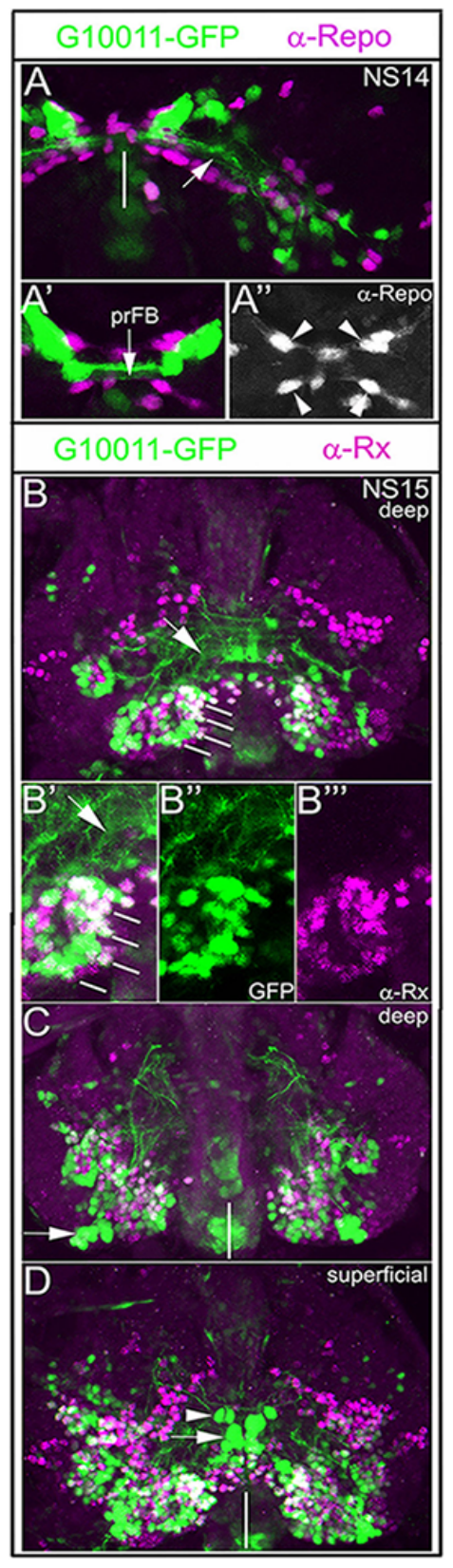

**Figure S 3:** Embryonic G10011 GFP-positive neurons establish the FB primordium (prFB). (**A-A’’**) Double-immuno-staining with α-GFP (green) and α-Repo antibody (magenta). Cell membranes of midline-associated glia (arrowheads in **A’’**) form a channel through which the DM1-DM4 progeny project their trajectories which constitute the prFB (arrows in **A** and **A’**; white line in **A** marks the midline). (**B-D**) Double-immuno-staining with α-GFP (green) and α-Rx antibody (magenta). (**B**) The prFB (arrow) is established by the progeny of the DM1-DM4 neuroblasts (white lines) (Andrade et al., 2019; Farnworth et al., 2020). (**B’-B’’’**) Many prFB neurons have been shown to express Rx protein (Farnworth et al., 2020). A subset of Rx-positive neurons co-expresses G10011-GFP. Arrow in **B’** indicates the trajectories of the four neuroblast progeny. (**C**) We interpret a group of posterior lateral located cells as AOTU neurons (arrow). (**D**). We interpret very large G10011-GFP-positive cells near the midline (white line) as neurosecretory cells of the prospective PI (arrow) and the “huging-expressing” cells (arrowhead), respectively.

**Figure S 4**

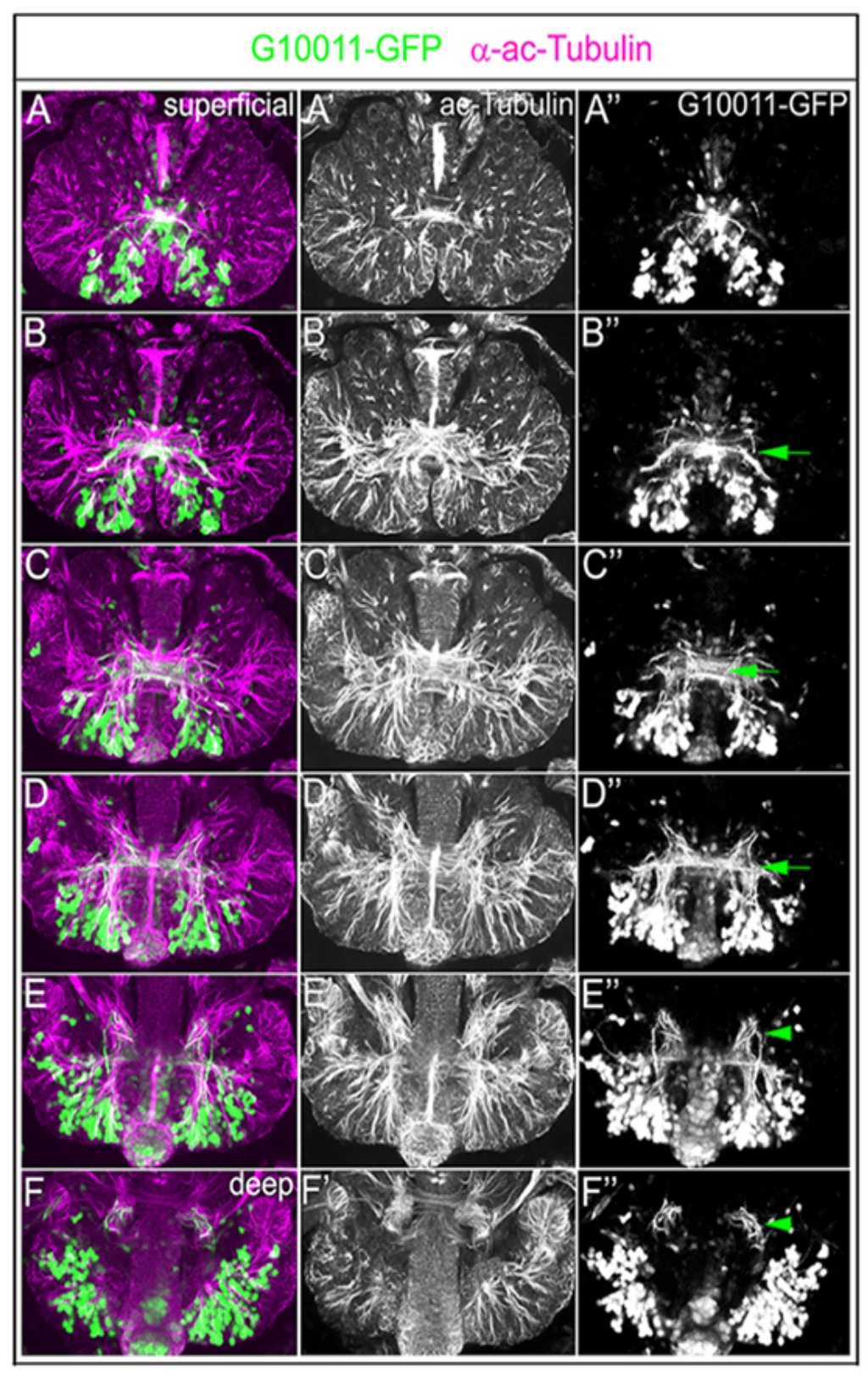

**Figure S 4:** G10011-GFP positive fascicles contribute to several major axon tracts in the embryonic brain. **(A-F)** G10011 brain at late stage NS15 stained with α-GFP (green) and α-acetylated Tubulin (magenta). Serial confocal sections were combined and visualized as maximum intensity projections to display individual anatomical features. (**A’-F’**) acetylated Tubulin only. (**A’’-F’’**) GFP only. (**B’’-D’’**) GFP-positive axon trajectories make multiple contributions to the commissural system (arrows). (**E’’,F’’**) GFP-positive fibres contribute to longitudinal axon tracts which extend towards the VNC (arrowheads).

**Figure S 5**

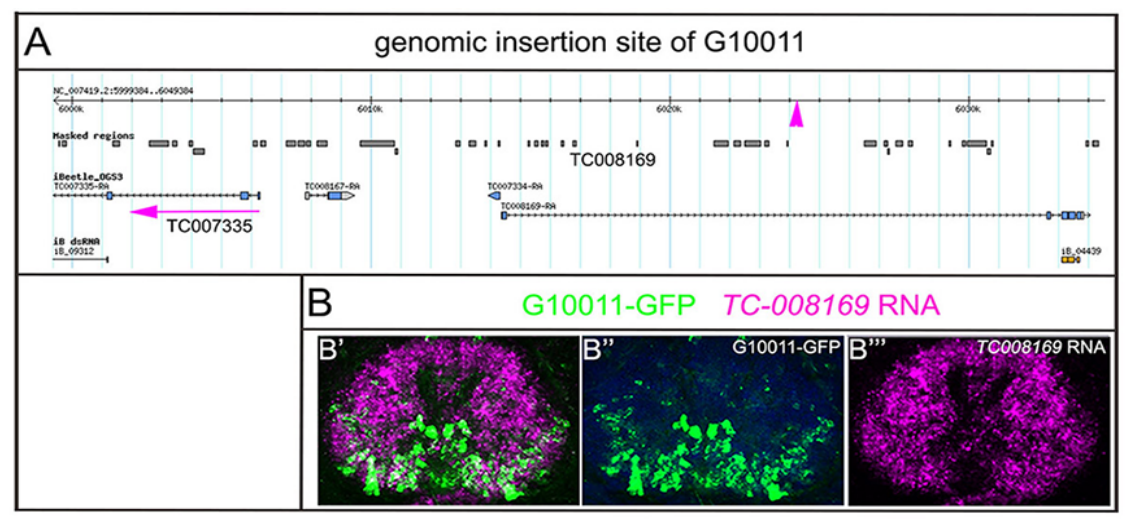

**Figure S 5: (A)** Genomic localization of the G10011 plasmid insertion site. The plasmid insertion was mapped to the position 602477 on the fourth chromosome (genome release 3.0). The insertion site is located in the first intron of the predicted gene *TC008169* (magenta arrowhead). We found no experimental evidence for the expression of the first *TC008169* exon suggesting that the plasmid insertion site is inter- rather than intragenic. *TC008169* encodes an EF1 hand protein. (**B**) To determine whether the expression of G10011-GFP reflects the expression of *TC008169*, we performed RNA in situ with a *TC008169* probe (magenta) combined with a α-GFP staining (green). **(B”’**) *TC008169* expression is pan-neural in the embryonic brain while G10011-GFP is expressed only in a subset of cells (**B’’**). We conclude that G10011-GFP is unlikely to reflect the expression of *TC008169*. The plasmid insertion site is located 18.5 kb upstream of the predicted gene *TC007335* (transcription start site 6006266, red arrow in (**A**)). The *TC007335* RNA in situ signal and the G10011-*GFP* RNA in situ signal co-localize in the embryonic and the adult brain indicating that G10011-GFP is a faithful reporter of *TC007335* RNA expression (refer to Figure 6).

**Figure S 6**

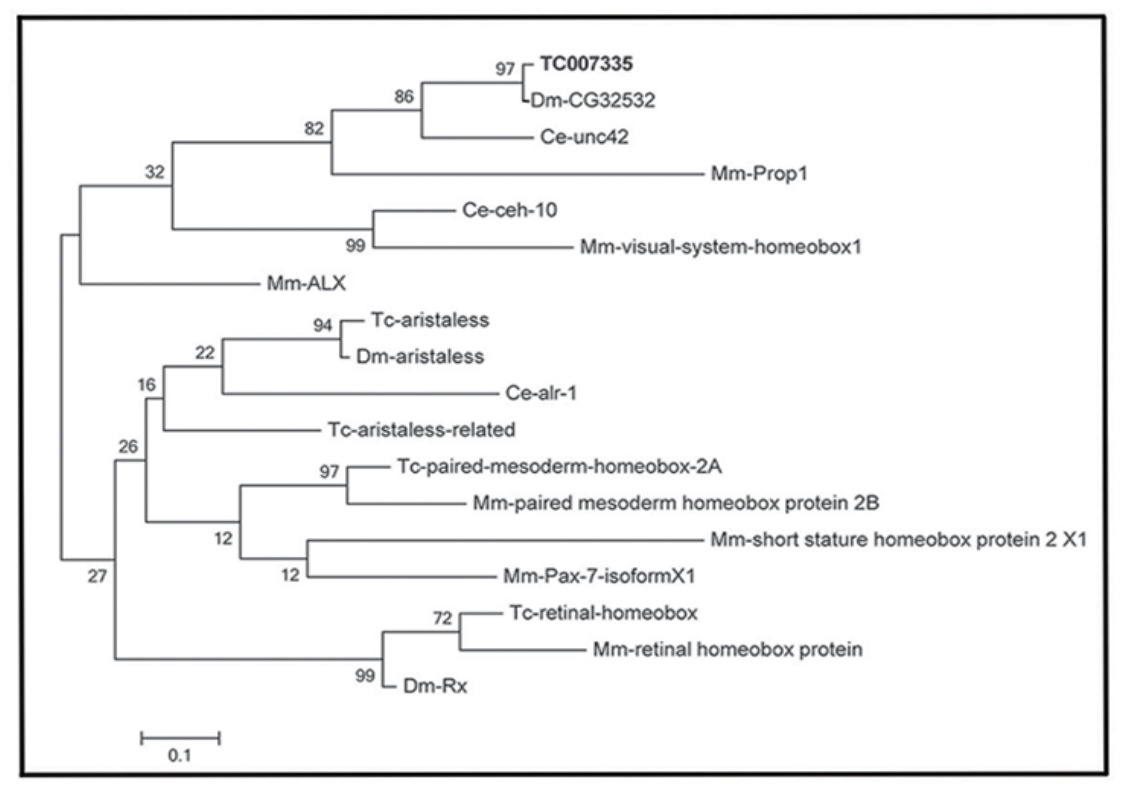

**Figure S6:** Phylogenetic tree reveals *D. melanogaster CG32532*, *C. elegans unc42* and *M. musculus Prop1* as single orthologs of *TC007335*. The TC007335 protein sequence was used to search the NCBI Ref-Seq databases for these species for the most similar proteins using blastp. Alignment was done using the Muscle algorithm as implemented in MEGA 6 (Tamura et al., 2013). The alignment was trimmed to remove all sequences with unclear alignment or gaps. We used Maximum Likelihood, UPGMA and neighbor joining algorithms as implemented in MEGA 6 with bootstrapping based on 500 replications to construct the phylogenetic tree. With all algorithms, the same orthology group is found (shown is the maximum likelihood tree with bootstrap values based on 500 replicates).

Original image stacks are available at

<https://figshare.com/projects/Additional_data_for_Garc_a_P_rez_et_al_Tribolium_shaking_hands_is_a_putative_terminal_selector_and_controls_axon_outgrowth_of_central_complex_neurons/93149>
